# Mapping the genetic architecture of human traits to cell types in the kidney identifies mechanisms of disease and potential treatments

**DOI:** 10.1101/2020.11.09.375592

**Authors:** Xin Sheng, Ziyuan Ma, Junnan Wu, Hongbo Liu, Chengxiang Qiu, Zhen Miao, Matthew J. Seasock, Matthew Palmer, Myung K. Shin, Kevin L. Duffin, Steven S. Pullen, Todd L. Edwards, Jacklyn N. Hellwege, Adriana M. Hung, Mingyao Li, Benjamin Voight, Thomas Coffman, Christopher D. Brown, Katalin Susztak

## Abstract

The functional interpretation of GWAS remains challenging due to cell-type dependent influences of genetic variants.

Here, we generated comprehensive maps of expression quantitative trait loci (eQTL) for 659 microdissected human kidney samples and identified cell-type eQTLs by mapping interactions between cell type abundance and genotype. Separately, we generated single cell open chromatin maps (by snATAC-seq) for human kidney samples. We highlight critical enrichment of proximal tubules in kidney function and endothelial cells and distal tubule segments in blood pressure by partitioning heritability using stratified LD-score regression to integrate GWAS with scRNA-seq and snATAC-seq data. Bayesian colocalization analysis nominated more than 200 genes for kidney function and hypertension. Our study clarifies the mechanism of the most commonly used antihypertensive and renal protective drugs and identifies drug repurposing opportunities for kidney disease.

**One Sentence Summary:** We define causal cell types, genes and mechanism for kidney dysfunction.

## Main Text

Chronic kidney disease (CKD) affects more than 800 million people worldwide (*1*). CKD causes toxic mineral, metabolite, salt, water accumulation (*2*), in addition to anemia, bone disease and hypertension leading to premature death (*3*). Therapeutic options for CKD have been limited. Guidelines mandate the use of agents blocking the renin-angiotensin-aldosterone-system (RAAS); such as angiotensin converting enzyme inhibitors (ACEi), and angiotensin receptor blockers (ARBs). Despite ACEi and ARBs were introduced more than 30 years ago, their mechanisms of action have not been fully clarified. As kidney disease is still responsible for 1 in 60 deaths worldwide, new therapeutics are desperately needed for CKD. Nelson et al. concluded, that pipeline drug targets with human genetic evidence of disease association are twice as likely to lead to approval (*4*), highlighting the critical importance of understanding disease genetics.

Kidney function is highly heritable, with estimates of 30-50% (*5*). Large population-based genome wide association studies (GWAS) have identified around 300 loci with statistically significant reproducible association with kidney function (eGFR; estimated glomerular filtration rate) (*5*). Despite the remarkable success of genetic signal mapping in GWAS, the functional interpretation of GWAS remains challenging. First, it is unclear in which tissues and cell types these variants active, and how they disrupt specific biological networks to impact disease risk. Second, more than 90% of the identified variants are located in the non-coding regions, complicating the precise identification of CKD risk target genes. Furthermore, extensive linkage disequilibrium (LD) in the human genome confounds efforts to pinpoint causal variants.

A commonly used approach to map variants to causal genes utilizes expression quantitative trait loci (eQTLs) (*6, 7*). The Genotype-Tissue Expression (GTEx) project has been a highly valuable resource to catalogue eQTLs across multiple human tissues, including kidney cortex with a limited sample size (n=73) (*6, 8*). Our and other groups have generated bulk glomerular and tubule specific eQTL data for about 120 kidney samples. This dataset was able to prioritize disease-causing genes for close to 24 of the GWAS identified loci (*7*). As both GWAS and eQTL have significant limitations on defining causal variants due to the linkage disequilibrium (*9*), the integration of these datasets focused around causal gene (rather causal variant) identification (*10, 11*). Functional genomics or epigenetics can provide an orthogonal information for GWAS interpretation by prioritizing likely causal variants (*12, 13*). Functionally important gene regulatory regions are mostly nucleosome-free and can be identified by transposase insertion (ATAC), DNAse hypersensitivity (*14*) and histone tail modifications (*15*). Datasets generated from whole human kidney tissue highlighted a modest enrichment of the GWAS signals in kidney specific enhancers (*12, 13*).

Another critical limitation of the eQTL and open chromatin analysis has been the cell type heterogeneity of the kidney. Recent single cell RNA-seq (scRNA-seq) studies have highlighted at least 30 different cell types in the kidney (*16, 17*). Studies analyzing a variety of blood cells identified (often rare) single disease driving cell types that was hard to read out using bulk datasets (*18*). Few approaches have been developed to identify disease-causal cell types. For example, cell-type-specific eQTLs have been generated using sorted (bulk) cell types (*19, 20*), an approach that is feasible for immune cells. It has been hard to generate cell-type specific eQTL data using scRNA-seq, as most current scRNA-seq methods suffer from uneven cell capture efficiency and low accuracy to define gene expression (*16*). Bulk tissue analysis coupled with computational cellular deconvolution have recently been developed for *in silico* estimation of cell type eQTLs, however, the accuracy of the computational methods has not been validated (*21-25*). Single cell epigenome studies, such as single-nucleus ATAC-seq (snATAC-seq), could provide potential insights into causal variants identification by overlapping GWAS variants with cell-type specific open chromatin regions (*26-28*).

Here, we combined several orthogonal approaches to prioritize causal cell types, variants, and genes for complex kidney disease-associated phenotypes. First, we identified expression of quantitative trait loci for 659 kidney samples by bulk eQTL analysis. We developed cell fraction adjusted SNP-cell type interaction (eQTL(ci)) analysis using estimated cell fractions, using *in silico* deconvolution (*29*). Separately, we generated single nuclear ATAC-seq (snATAC-seq) maps for the human kidney and assessed enrichment of GWAS heritability in each cell type, obtained from scRNA-seq and snATAC-seq data, using stratified LD-score regression. Our study not only highlighted critical cell type convergence of kidney associated traits, but also nominated causal variants, cell types and regulatory mechanisms for renal disease and defines mechanisms for commonly used antihypertensive and renoprotective drugs.

## Results

### Cell fraction adjusted eQTL analysis of microdissected human kidney samples

While GWAS efficiently describe the genotype association with specific traits and diseases, understanding the genotype effect on gene expression (eQTL) has been essential to nominate target genes for GWAS variants. Prior eQTL studies have been limited, as they used whole organ tissue, comprised of many cell types, and small sample sizes (*7, 30*). Here we collected a large number of human kidneys from subjects with different ethnicities, disease status and disease stages (**Tables S1 and S2**). To reduce cell heterogeneity, we manually microdissected each tissue sample into glomerular and tubule compartments, and performed RNA-sequencing and genotyping on 659 samples (356 tubules and 303 glomeruli) (Fig. 1A). We initially performed eQTL analysis separately in glomerular and tubule samples following the GTEx v7 pipeline (*6*). The analysis took age, gender, collection site, RNA integrity (RIN), sequencing parameters (batch effect, read depth, and sequencing types), genotyping principle components (PCs), and kidney disease severity into consideration. This first cis-eQTL analysis identified 3,599 and 5,871 eGenes (genes that showed association with at least one SNP at a q-value<0.05) in tubule and glomerular samples, respectively (**Fig. S1A and B**).

**Figure 1.**
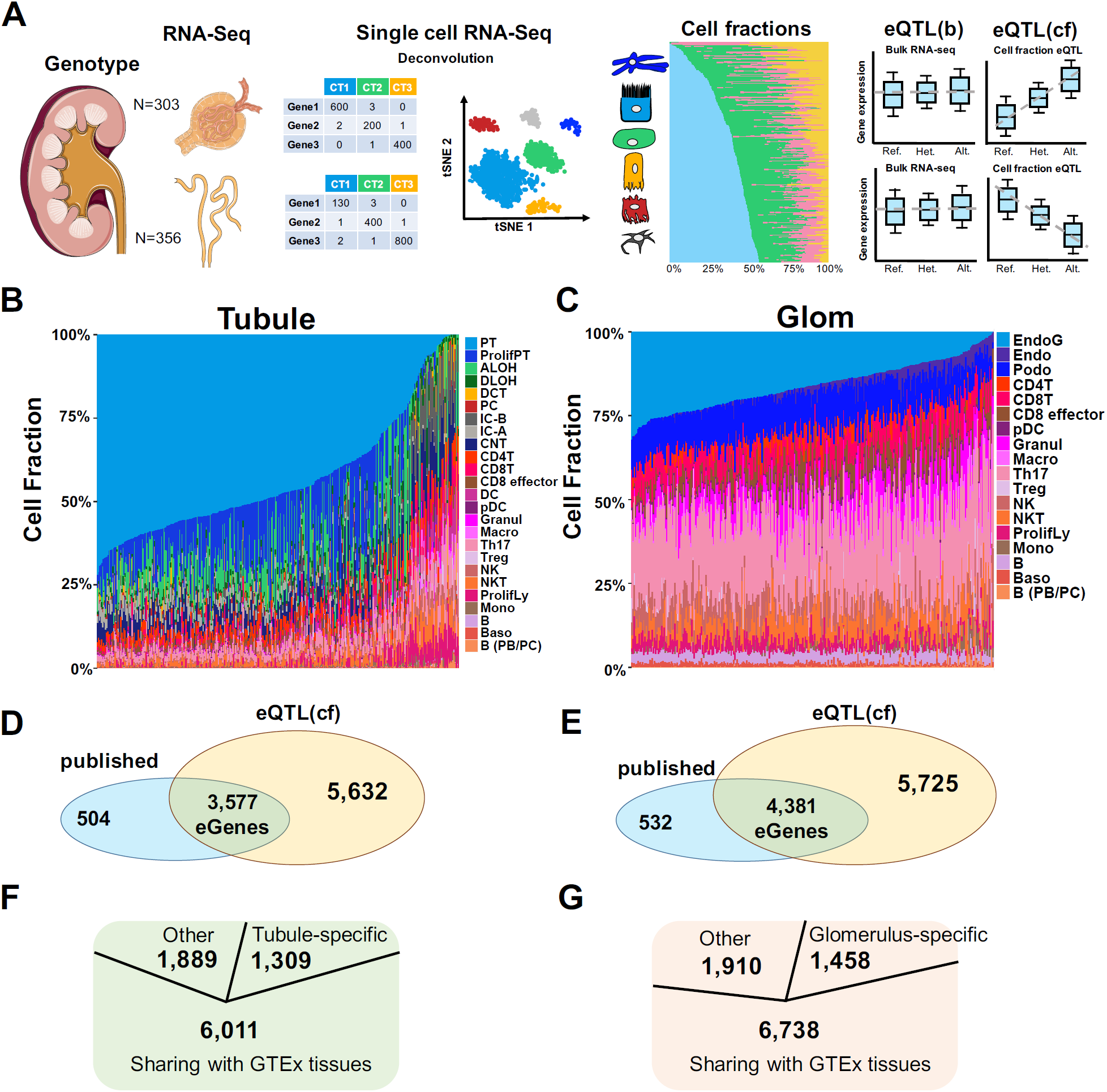
Cell fraction adjusted expression quantitative trait analysis of microdissected human kidney samples. A. Experimental scheme. We collected high quality genotype and RNA-seq data for 303 human microdissected kidney glomeruli and 359 tubule samples. Cell fractions were estimated by deconvolution method using scRNA-seq data (*29*) as the reference. eQTLs were then identified using bulk and cell fraction adjusted models. B. The distribution of the relative cell fractions across 359 microdissected human kidney tubule samples. PT: proximal tubule, ProlifPT: proliferating PT cells, ALOH: ascending loop of Henle, DLOH: descending loop of Henle, DCT: distal convoluted tubule, PC: collecting duct principal cells, IC-B: beta intercalated cells, IC-A: alpha intercalated cells, CNT: connecting tubule, CD4T: CD4 T cells, CD8T: CD8 T cells, CD8 effector: CD8 effector cells, DC: CD11b+ dendritic cells, pDC: plasmacytoid DC, Granul: granulocyte, Macro: macrophage, Th17: T helper 17 cells, Treg: regulatory T cells, NK: natural killer cells, NKT: natural killer T cells, ProlifLy: proliferating lymphocyte cells, Mono: monocyte, B: B lymphocyte, Baso: Basophil, B (PB/PC): memory B cells from the peripheral blood or plasma cells. C. The distribution of the relative cell fractions across 303 microdissected human kidney glomerular samples. EndoG: glomerular endothelial cells, Endo: endothelial cells, Podo: podocyte, CD4T: CD4 T cells, CD8T: CD8 T cells, CD 8effector: CD8 effector cells, pDC: plasmacytoid DC, Granul: granulocyte, Macro: macrophage, Th17: T helper 17 cells, Treg: regulatory T cells, NK: natural killer cells, NKT: natural killer T cells, ProlifLy: proliferating lymphocyte cells, Mono: monocyte, B: B lymphocyte, Baso: Basophil, B (PB/PC): memory B cells from the peripheral blood or plasma cells. D. Venn diagram illustrating the direct overlap of previously published (*7*) and newly identified significant eGenes in microdissected human kidney tubules E. Venn diagram illustrating the direct overlap of previously published (*7*) and newly identified significant eGenes in microdissected human kidney glomeruli F. Diagram illustrating the number of tubule-specific eGenes identified by meta-analysis of eQTLs from 49 human tissues (48 GTEx v7 (*34*) tissues plus tubule). G. Diagram illustrating the number of glomerulus-specific eGenes identified by meta-analysis of eQTLs from 49 human tissues (48 GTEx v7 (*34*) tissues plus glomerulus).

Since we observed heterogeneity in the relative distributions of cell populations across the RNA-seq samples (Fig. 1B **and** C), we hypothesized that considering the cell population distributions of each sample would improve the quality of eQTL inferences by reducing heterogeneity and increasing our power to detect associations within tissues and/or cell types. We used kidney single cell gene expression matrix (*29*) and an *in silico* deconvolution method implemented in CIBERSORTx (*22, 25*) to estimate cell fractions in each microdissected sample (Fig. 1A, **Methods**). As expected, we observed that proximal tubules (PT) represented the major cell type in the human kidney tubule samples (Fig. 1B) and endothelial cells in glomeruli (Fig. 1C). Next, we performed a second eQTL analysis by further adjusting for cell population differences across samples. We found that comparing to our initial eQTL analysis model without considering the cell fractions, the inclusion of cell fractions has improved the eGene identification by 28% (1,256 genes) and 18% (1,194 genes) in tubule (**Fig. S1A**) and glomerular (**Fig. S1B**) compartments, respectively.

Our final eQTL(cf) model included estimated latent variable adjustment by PEER factors in addition to cell fraction adjustment (**Methods**). This analysis identified 9,209 and 10,106 eGenes (Fig. 1D **and** E), including 6,821 and 7,501 protein coding genes, and 865,410 and 897,548 significant SNP-gene pairs (at a q-value <0.05) (*31*), in tubular and glomeruli samples, respectively. It is important to note that this eQTL(cf) model replicated ∼90% eGenes identified by the general eQTL model (only including PEER factors) (**Fig. S1C and D**), which is consistent with previous findings that PEER factors modeled cellular heterogeneity reasonably well (**Fig. S2**) (*32*).

Our eQTL(cf) model identified more than twice as many eGenes as reported in our previous publication (*7*) (**Fig. S3**), which is likely due to increasing power of eGene discovery with a larger sample size and better adjustments for confounders. Several genes, such as *KIAA1841* (**Fig. S4A**) and *MRPL43* (**Fig. S4B**) did not reach significance in the previous eQTL analysis, but showed strong associations between gene expression and genotypes in our eQTL(cf) model, illustrating the power of the new dataset. The significant SNP-gene pairs identified by our previous study could be successfully replicated by the eQTL(cf) model with **π**_1_=0.95 in tubule and **π**_1_=0.99 in glomeruli, respectively (*33*). There was a 90% direct overlap between the newly and previously reported eGene list (Fig. 1D **and** E). Meta-analysis combining evidence from results with 48 tissues in the published GTEx data showed a large number of shared eGenes (*34*) and also 1,309 tubule-specific (Fig. 1F) and 1,458 glomerulus-specific (Fig. 1G) eGenes (M-value>0.9) in our eQTL(cf) study.

In summary, here we report bulk and cell fraction adjusted eQTL data for more than 600 microdissected human kidney glomeruli and tubule samples. The increased sample size and improved cell fraction estimation have more than doubled the number of identified eGenes and revealed a large number of kidney compartment specific eGenes.

### Defining cell-type dependent activities of genetic variants on gene expression

In order to identify eQTLs that modulate gene expression in a cell-type specific manner, we adopted a linear regression model that included an interaction term between cell fraction and genotype (**Methods**). We refer to these eQTLs as cell type interacting eQTLs (eQTL(ci)) (*8*) (Fig. 2A). For example, kidneys of subjects with higher allele dosage of G (SNP rs12481710), showed a positive correlation between PT fractions and *SYCP2* (Synaptonemal complex protein 2) expression (Fig. 2B), indicating a cell-type specific genotype-gene expression interaction. In total, we identified 1,613 and 713 protein-coding eQTL(ci) genes in tubule (**Table S3**) and glomerular (**Table S4**) samples, respectively (Fig. 2C) (*8*), by evaluating the significance of the interaction effect between genotype and cell fraction (**Methods**) in 23 cell types (n=13 in tubule, n=10 in glomeruli) of the kidney. Kidney PT cells showed the highest number of eQTL(ci)s in the microdissected human kidney tubule compartment (Fig. 2C), while glomerular endothelial cells (EndoG) showed the highest number of eQTL(ci)s in glomeruli (Fig. 2C). The number of eQTL(ci)s for each cell type and sample size was in line with the previous publication (*8*).

**Figure 2.**
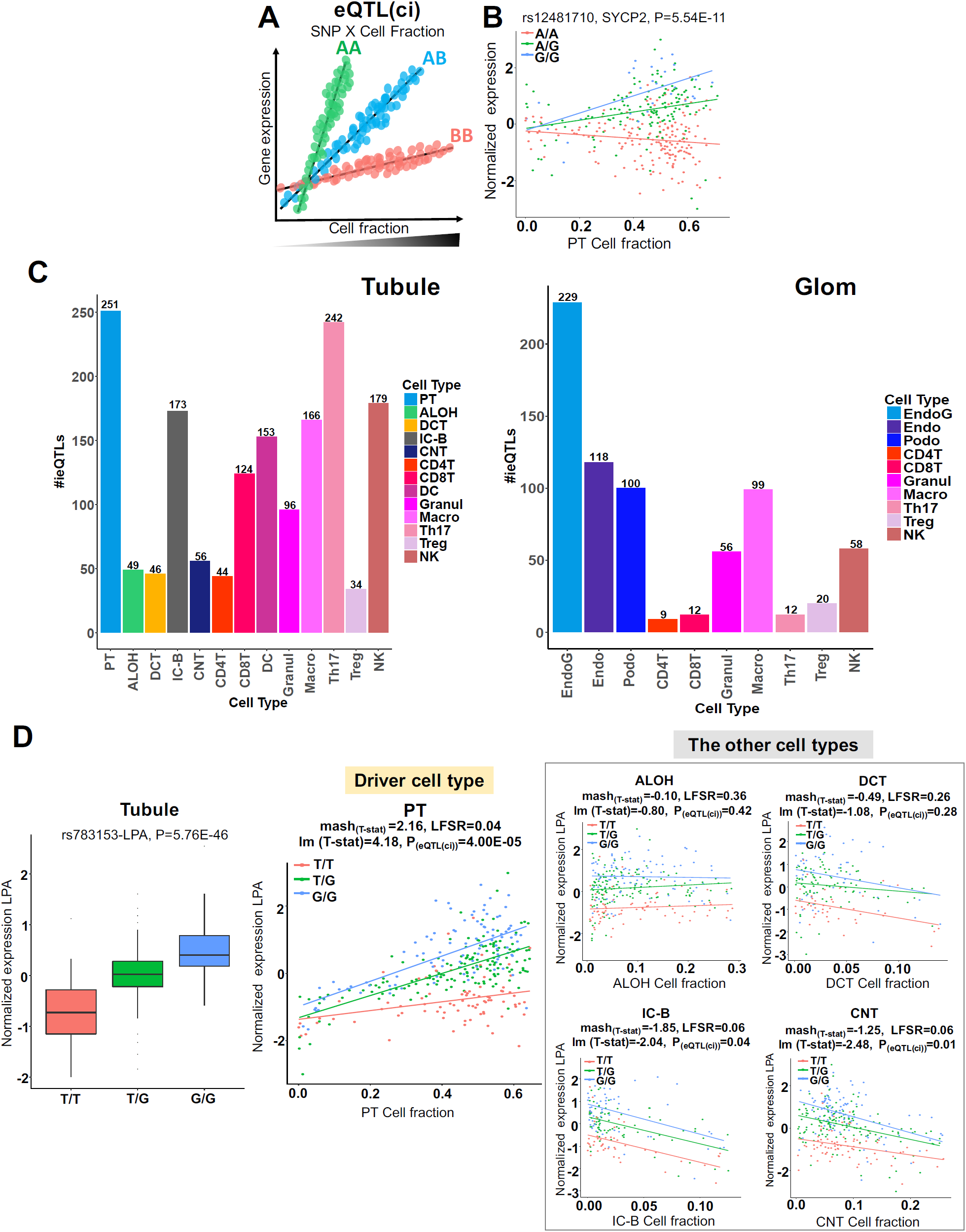
Defining cell-type dependent activities of genetic variants on gene expression. A. Scheme of cell type interaction eQTL (eQTL(ci)). For a specific SNP, the correlation between gene expression variations and cell fraction changes are significantly varying across individuals with different genotypes on this locus. B. The y-axis is the relative expression of *SYCP2* in human kidney tubules (N=356 samples), the x-axis is the PT cell fraction colored by subject (rs12481710) genotype A/A (red), A/G (green) and G/G (blue). P value was calculated by eQTL(ci) model. Each data point represents a single subject. The y-axis is the normalized expression of *SYCP2* in human tubules (N=356 samples), the x-axis is the cell fraction colored by subject genotype A/A (red), A/G (green) and G/G (blue) at the rs12481710 locus. C. The number of identified eQTL(ci)s in each cell types in human kidney tubules (left panel) and glomeruli (right panel). EndoG: glomerular endothelial cells, Endo: endothelial cells, Podo: podocyte, PT: proximal tubule, ALOH: ascending loop of Henle, DCT: distal convoluted tubule, IC-B: beta intercalated cells, CNT: connecting tubule, CD4T: CD4 T cells, CD8T: CD8 T cells, DC: CD11b+ dendritic cells, Granul: granulocyte, Macro: macrophage, Th17: T helper 17 cells, Treg: regulatory T cells, NK: natural killer cells. D. A PT specific eQTL(ci) for *LPA*. The leftmost panel illustrates the association between genotype of SNP rs783153 and gene expression of *LPA* in bulk tubule samples (N=356 samples). The y-axis is the normalized expression of *LPA* in human kidney tubules, the x-axis is the genotype at rs783153 locus. Center lines show the medians; box limits indicate the 25^th^ and 75^th^ percentiles; whiskers extend to the 5^th^ and 95^th^ percentiles; outliers are represented by dots. P was calculated by linear regression eQTL(cf) model. The following panels show PT cell-type-specificity of this eQTL(ci) across 5 different kidney function cell types in human tubules (N=356 samples). The y-axis is the normalized expression of *LPA* in human tubules (N=356 samples), the x-axis is the cell fraction colored by subject genotype T/T (red), T/G (green) and G/G (blue) at the rs783153 locus. lm (T-stat): T-statistics calculated from linear model of eQTL(ci), mash_(T-stat)_: T-statisitcs estimated by *mash*. PT: proximal tubule, ALOH: ascending loop of Henle, DCT: distal convoluted tubule, IC-B: beta intercalated cells, CNT: connecting tubule. Each data point represents one sample.

To characterize the cell type specificity of eQTL(ci)s, we examined the eQTL(ci) sharing across kidney cell types using a multivariate adaptive shrinkage method (*mash*) (*35*), which improves the effect size estimation for the cell fraction and genotype interaction. *Mash* leverages the correlations of effect sizes across all cell types to re-estimate eQTL(ci) effect size at each locus. Cell type specific eQTL(ci) was defined as the eQTL(ci) at LFSR<0.05 (the local false sign rate (*36*)) only in a single cell type. We found that using this stringent threshold, about 18% (n=3,544 of 20,146 SNP-Gene pairs) eQTL(ci)s were cell type specific across the 23 highly correlated kidney and immune cell types (**Fig. S5**).

Here, we show that lipoprotein A (*LPA*) expression significantly correlated with rs783153 (eQTL(cf) P-value=5.76E-46) in tubule (Fig 2D). This SNP modified the relationship between PT cell fraction and *LPA* expression, such as *LPA* expression in subjects with increasing dosage of G allele is increased with increasing PT fractions. The *LPA* gene expression showed a greater increase with PT fractions in individuals with genotype G/G at rs783153, compared to individuals with genotype T/G and T/T, which is consistent with the bulk eQTL data. T-statistics for eQTL(ci)s in each cell type of the kidney tubules, which were estimated using *mash* and our eQTL(ci) linear regression model, show consistent regulatory directions in each cell type. Interestingly, the eQTL effect observed in tubule was consistent with the PT specific regulatory eQTL(ci) effect (Fig 2D) (with LFSR<0.05 only in PT cells). These results indicate that for this eQTL pair (rs783153-*LPA*), the PT eQTL(ci) acted as the major driver of eQTL(cf) effect in the bulk tubule tissue. Similarly, we found that the eQTL pair (rs8023408-*SV2B* (Synaptic Vesicle Glycoprotein 2B)), which showed that individuals with genotype A/A had higher *SV2B* expression in the bulk glomerular tissue (**Fig S7**), was related to a EndoG specific eQTL(ci) effect. There was a greater increase in *SV2B* expression as the fraction of EndoG increased in subjects with A/A genotype. This was an EndoG specific eQTL(ci) (LFSR<0.05 only in EndoG cells) and the genotype did not significantly modify the cell fraction and gene expression relationship in other cell types. This indicated that the EndoG specific eQTL(ci) (rs783153-*LPA*) effect was the key driver of the eQTL(cf) (**Fig. S6**) in the bulk glomerular tissue. Taken together, using the eQTL(ci) model, we have defined a list of genotype-gene expression correlations that is likely driven by a specific cell type in the human kidney.

### Single cell resolution regulatory maps for the human kidney

Cell-type specific regulatory elements have not been mapped for the human kidney, which is an important knowledge gap as functionally relevant GWAS variants are likely enriched in cell type specific regulatory regions (*28*). To characterize gene regulatory regions at a cellular resolution, we generated single nuclei accessible chromatin information for 12,720 human kidney cells using the 10x Genomics snATAC-seq platform (Fig. 3A). After initial quality control, we identified 8 distinct cell clusters (Fig. 3B). By comparing the read density of the promoter regions in each cluster, we found that the initial clustering showed a clear match with previously identified key kidney gene expression markers (*16, 37*), such as *KDR* for endothelial cells (Endo), *NPHS2* for podocytes (Podo), *LRP2* for proximal tubule cells (PT), *UMOD* for loop of Henle (LOH), *SLC12A3* for distal convoluted tubule (DCT), *AQP2* for principal cell (PC), *ATP6V1G3* for intercalated cells (IC) of the collecting duct (CD) and *LST1* for immune cells (Immune), respectively (Fig. 3C **and Fig. S7**), indicating the robustness of the independently identified clusters. This analysis identified a total of 359,019 accessible chromatin peaks in the entire dataset for the human kidney (**Methods**). Single cell open chromatin peaks were enriched in human kidney ChromHMM defined promoter and enhancer regions (**Fig. S8**). For example, 38% of open chromatin regions observed in PT cells overlapped with enhancer regions and 17% of peaks overlapped with promoter regions in adult bulk human kidney samples (**Table S5**).

**Figure 3.**
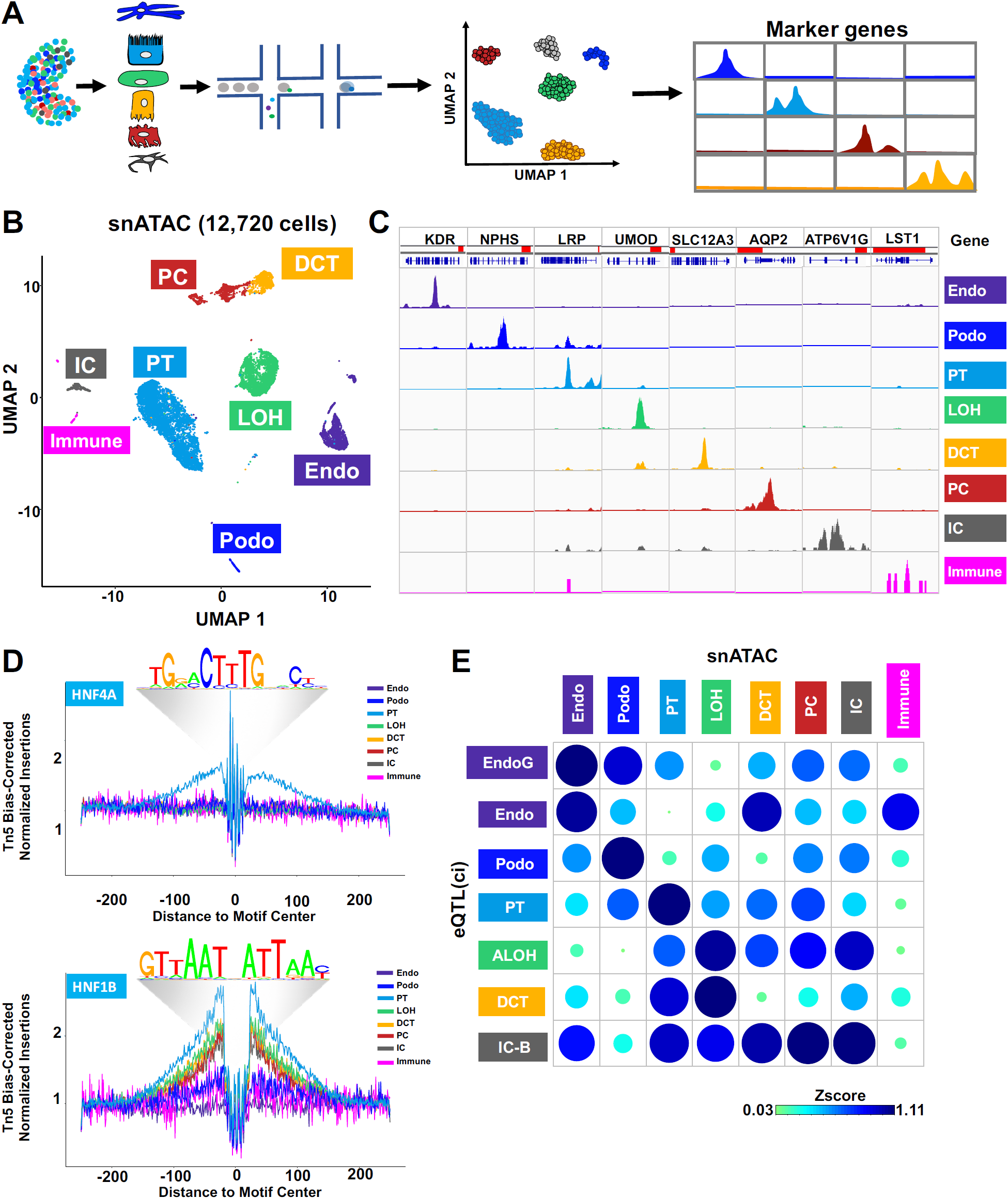
Single cell resolution regulatory maps for the human kidney. A. Experimental scheme of snATAC-seq. Single nuclei accessible chromatin information was generated for 12,720 human kidney cells, using the 10X Genomics snATAC-seq platform. Kidney cell clusters were then identified and open chromatin peaks were called for each cell cluster. B. The UMAP of human kidney snATAC-seq data. 8 distinct cell clusters were identified using previously defined cell type marker genes (*16*). Endo: endothelial cells, Podo: podocyte, PT: proximal tubule, LOH: loop of Henle, DCT: distal convoluted tubule, PC: collecting duct principal cells, IC: collecting duct intercalated cells, Immune: immune cells. C. Genome browser plots showing aggregate read density for cells within each cell cluster in the promoter region of each cell type marker gene: *KDR* (endothelial cells), *NPHS2* (podocyte), *LRP2* (proximal tubule), *UMOD* (LOH), *SLC12A3* (distal convoluted tubule), *AQP2* (collecting duct principal cells), *ATP6V1G3* (collecting duct intercalated cells) and *LST1* (immune cells). The promoter region for each gene is highlighted in red color. D. Footprinting analysis of the HNF4A (top panel) and HNF1B (bottom panel) transcription factors across 8 major cell types. The motif logos are shown above. E. Bubble plots of enrichment of eQTL(ci)s for each cell type (row) and open human kidney snATAC-seq chromatin regions of each cell cluster (Column). The size and the color of the bubble indicates enrichment (Z-score). EndoG: glomerular endothelial cells, Endo: endothelial cells, Podo: podocyte, PT: proximal tubule, LOH: loop of Henle, ALOH: ascending loop of Henle, DCT: distal convoluted tubule, PC: collecting duct principal cells, IC: collecting duct intercalated cells, IC-B: beta intercalated cells, Immune: immune cells.

Next, we generated a list of differentially accessible chromatin regions (DARs) by comparing open chromatin regions across clusters and identified 60,661 cell-type specific peaks (**Methods**). To identify transcription factors (TFs) that predominantly occupy cell type specific open chromatin regions, we performed motif enrichment analysis of differentially accessible regions (DARs) for each cell type using chromVAR (*38*) and Homer (*39*). This analysis identified HNF4A motif enrichment in proximal tubules, ESRRB in loop of Henle, and ERG in endothelial cells (*40*) (**Fig. S9**). Furthermore, TF footprinting analysis across the 8 major cell types (Fig. 3D **and Fig. S10**) showed cell type specific enrichment of TF binding to HNF4A (Fig. 3D) and HNF4G **(Fig. S10)** in proximal tubule, and tubule compartment specific motif enrichment of TF HNF1B (Fig. 3D). Overall, these results provide a reference landscape for chromatin accessibility in the adult human kidneys at single cell resolution.

The cell-type specificity of the eQTL(ci) has not been experimentally validated yet. We next estimated the enrichment of eQTL(ci) variants in the associated cell type specific open chromatin regions (by snATAC-seq) (*41*) (**Methods**). We found that for most common kidney cell types (EndoG, Endo, Podo, PT, ALOH and IC-B), eQTL(ci) SNPs showed the strongest enrichment in the open chromatin regions of the corresponding cell type (Fig. 3E), indicating consistency between cell type interaction eQTLs and the corresponding cell type specific regulatory regions. The similarity of open chromatin in DCT and LOH (**Fig. S9**) cells likely explained the overlap we observed for these cells.

To summarize, we generated a single cell resolution regulatory map of genetic regulation of gene expression for the human kidney. We found that our cell-type specific eQTL(ci)s enriched in open chromatin regions of the corresponding cell type using snATAC-seq data, indicating accuracy of the computational eQTL(ci)s identification.

### Single cell annotation highlights cell-type convergence of kidney relevant endophenotypes

To prioritize potential driver cell types for specific kidney dysfunction associated endophenotypes (such as glomerular filtration rate, albuminuria, and hypertension), we first calculated and ranked the expression specificity of each gene for each cell type from the single cell RNA-seq data (*16*). For a cell type, higher gene expression specificity deciles were associated with increasing levels of cell type expression specificity (**Methods**). Next, we assessed the partitioning heritability enrichment from GWAS summary statistics in genes of each decile (Fig. 4A) using linkage disequilibrium score regression (LDSC) (*42*). We observed that the enrichment of eGFR SNP heritability increased with increasing gene expression specificity for PT cells (Fig. 4B). Similar positive correlation was observed between SBP and DCT (Fig. 4C), PC (**Fig. S11**) and endothelial cells (Fig. 4D), suggesting the importance of these cell types for the tested endophenotypes.

**Figure 4.**
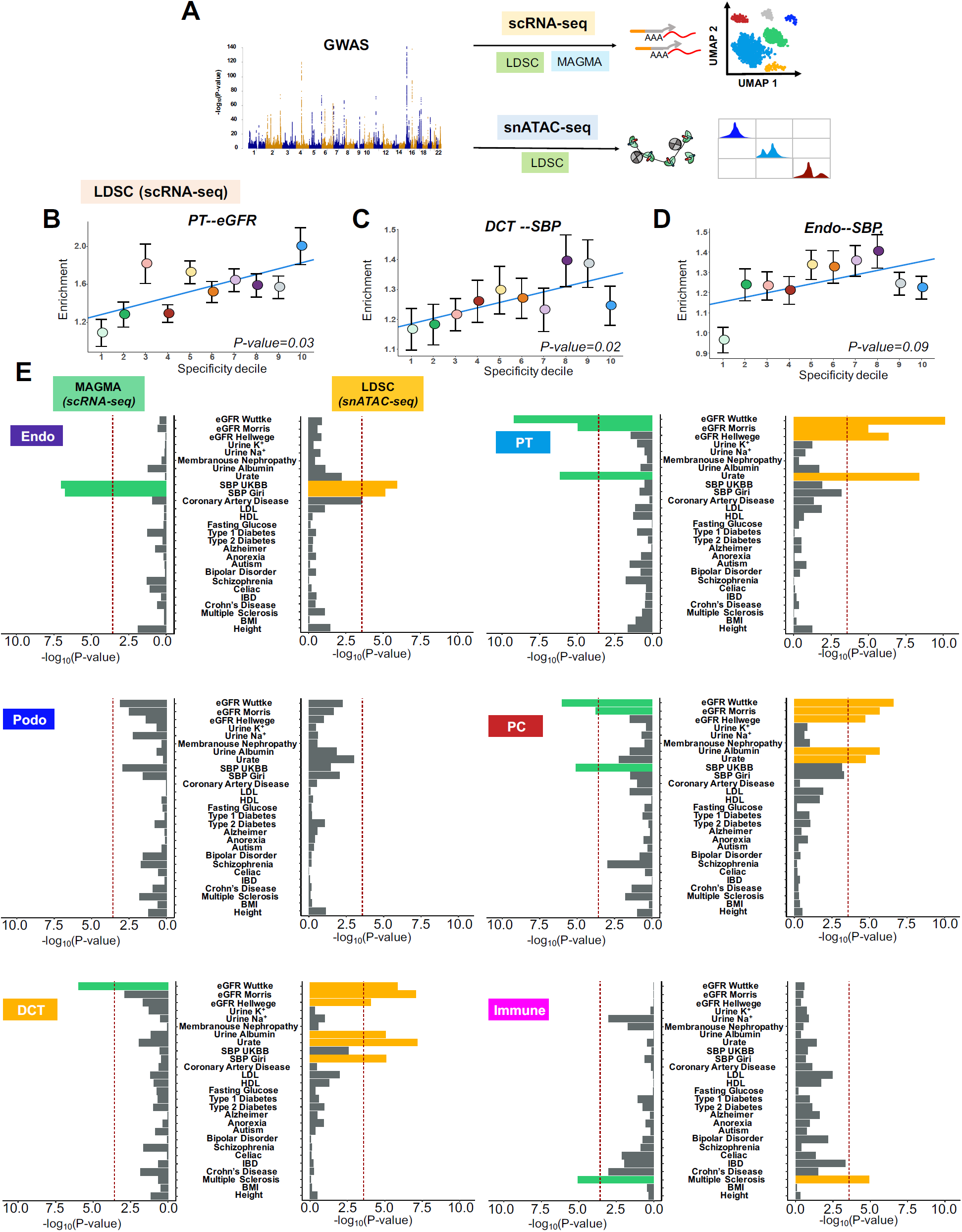
Single cell annotation highlights cell-type convergence of kidney relevant endophenotypes. A. Experimental scheme of cell type-specific GWAS trait heritability enrichment analysis. Here we applied MAGMA to the scRNA-seq data and LDSC to the scRNA-seq and snATAC-seq data to assess the GWAS per-SNP heritability enrichment. B. Enrichment of eGFR-SNP heritability in each of the specificity deciles for proximal tubule cells calculated using LDSC from kidney scRNA-seq data (*16*). C. Enrichment of SBP-SNP heritability in each of the specificity deciles in distal convoluted tubule (DCT) cells. D. Enrichment of SBP-SNP heritability in each of the specific deciles for endothelial cells. X-axis is the gene expression specificity decile, Y-axis is the enrichment value calculated by LDSC. Colors of dots represent different specificity deciles. Blue line shows the linear regression slope fitted to the enrichment values. Error bars indicate the 95% confidence intervals. E. Evaluation of enrichment of common GWAS variants in each kidney cell type scRNA-seq data (*16*) using MAGMA (the left panels) and human kidney snATAC-seq data using LDSC (the right panels). X-axis represents 24 different GWAS traits and y-axis is the –log_10_(P-value). P-value was estimated by MAGMA (the left panels) and LDSC (the right panels). Endo: endothelial cells, PT: proximal tubule, Podo: podocyte, PC: collecting duct principal cells, DCT: distal convoluted tubule, Immune: T lymphocyte.

Next, we adopted a generalized gene set enrichment analysis established in MAGMA (*43*) to evaluate whether gene-level genetic association with kidney relevant phenotypes increased linearly with cell type specificity (**Methods**). We have analyzed 24 different GWAS traits (Fig. 4E **and Fig. S12**) including three independent eGFR GWAS (*5, 44, 45*), two systolic blood pressure (SBP) GWAS (*46*), serum urate (*47*) (Urate), urinary albumin-to-creatinine ratio (UACR) (*48*), urinary sodium (Na^+^) and potassium (K^+^) excretion (*49*), and cardiometabolic traits. As established earlier, multiple sclerosis showed significant enrichment in lymphocytes also in our kidney dataset (*50*). We found the strongest enrichment for kidney function (eGFR) and serum urate level in proximal tubules (*51*) (Fig. 4E). Genetic signals for blood pressure showed enrichment in endothelial cells.

Simultaneously, we adopted a third independent analysis by implementing LDSC using the single cell open chromatin information. We note that snATAC-seq LDSC evaluates the heritability of GWAS variants under snATAC-seq peaks, which is distinct from the measured gene expression output of scRNA-seq. We found a highly consistent enrichment of multiple eGFR GWAS signals in PT cells. Serum urate signals were the most enriched in PT cells comparing to other cells (Figs. 4E **and Fig. S12**). We observed enrichment of SBP genetic signals in endothelial cells, tubule cells in the distal nephron (DCT and PC cells) (Fig. 4E **and Fig. S12**).

To summarize, using a variety of orthogonal datasets and methods, we observed important cell type convergence for several tested endophenotypes. The strongest of all was the enrichment between eGFR associated genetic signals and proximal tubule cells (Fig. 4B **and** E) and SBP associated genetic signals in endothelial (Fig. 4E) and DCT cells (Figs. 4C, E **and S11**).

### Comprehensive gene prioritization provides new mechanistic insights into kidney function and blood pressure regulation

Next, we prioritized genes that likely mediate phenotypic variations for specific renal endophenotypes. Using the Bayesian test for colocalization (implemented in coloc) (*52*) to integrate 6 kidney related traits including eGFR, SBP, serum urate, UACR, urinary Na^+^ and K^+^, and kidney microdissected compartment eQTL(cf)s data, we identified 240 colocalized protein coding genes in the tubule (**Table S6**) and 230 genes in the glomerular compartment (**Table S7**). At these regions, causal genetic variants driving gene expression changes and the variations of these 6 renal endophenotypes were shared. The analysis prioritized 182 genes for kidney function (eGFR) (Fig. 5A **and Table S8**) (*5*). This number was markedly increased compared to the 27 genes prioritized by earlier studies (*7*). Similar colocalization analysis prioritized 88 genes for the 340 GWAS loci (*46*) associated with SBP (Fig. 5B **and Table S9**), where the causal genetic variants associated with blood pressure and gene expression were shared. The distribution of the eGFR and blood pressure prioritized genes in the glomerular and tubular compartments across the genome is illustrated in Fig. 5C **and Fig. S13**. Only 10 genes were prioritized for both SBP and eGFR, indicating that genetic architecture of these two traits are likely distinct, which is consistent with the LDSC analysis estimating the genetic correlation of these 6 GWAS traits (**Fig. S14**) (*9*). One of the loci and genes that was associated with both traits was *FGF5* (Fibroblast Growth Factor 5). Allelic dosage of G at the SNP rs3733336 position was associated with a higher *FGF5* expression levels in glomeruli (**Fig. S15A**). There was a strong correlation between the eQTL and GWAS effect sizes of SNPs located within ±100 kb of this locus for both SBP (**Fig. S15B**) and eGFR GWAS (**Fig. S15C**).

**Figure 5.**
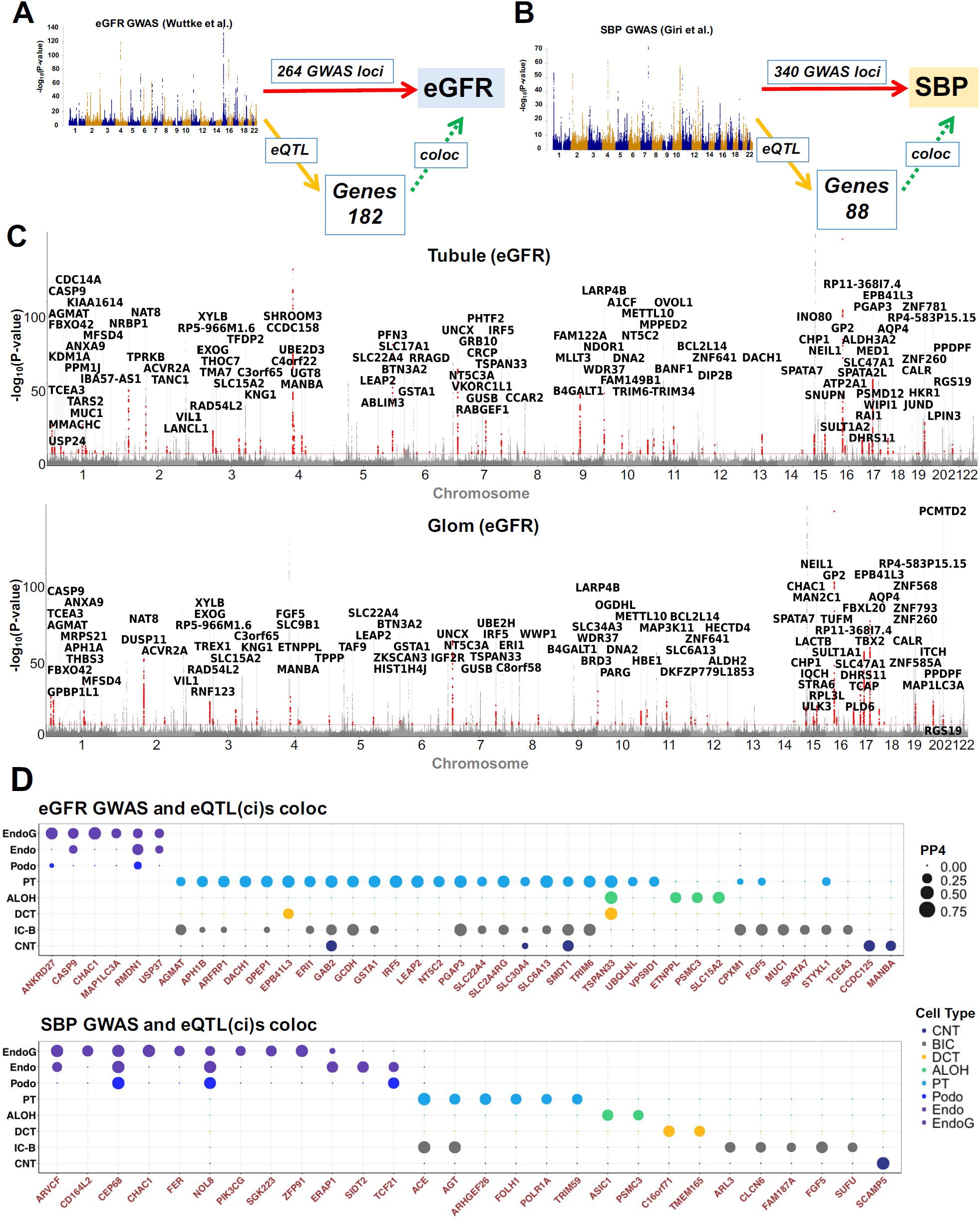
Comprehensive gene prioritization provides new mechanistic insights into kidney function and blood pressure regulation. A. Experimental scheme. We used Bayesian colocalization which combined information from eQTL(cf)s to annotate 264 eGFR associated loci (*5*) and prioritized 182 causal genes for kidney disease, where the causal variants for gene expression and kidney function are shared. B. Experimental scheme. We used Bayesian colocalization which combined information from eQTL(cf)s to annotate 340 SBP GWAS associated loci (*46*) and prioritized 88 causal genes for hypertension, where the causal variants for gene expression and HTN are shared. C. Miami plot showing coloc (*10*) prioritized genes for eGFR GWAS in tubules and glomeruli by eQTL(cf)s. Red dots define prioritized protein coding genes. X-axis represents the chromosome. Y-axis represents the GWAS significance –log_10_(P-value). D. Bubble plots showing coloc (*10*) prioritized cell type specific (eQTL(ci)) genes for eGFR and SBP GWAS loci. The size of the dot correlated with the posterior possibility of colocalization (PP4). EndoG: glomerular endothelial cells, Endo: endothelial cells, Podo: podocyte, PT: proximal tubule, ALOH: ascending loop of Henle, DCT: distal convoluted tubule, IC-B: beta intercalated cells, CNT: connecting tubule.

To understand whether the eQTL(ci) data could be used to improve GWAS target identification and also to nominate causal cell types, we repeated the Bayesian colocalization analysis using GWAS summary data of the 6 kidney related traits and the eQTL(ci)s data. We prioritized 61 potentially causal protein coding genes, where variants for the 6 renal endophenotypes and cell-type specific gene regulation were shared (**Table S10**). We again observed a significant enrichment of PT eQTL(ci)s and eGFR GWAS, such as 23 of the 40 showed colocalization with interaction PT eQTL(ci)s. For SBP GWAS, we observed significant enrichment for EndoG eQTL (ci)s, where 9 of the 28 identified variants showed cell type enrichment (Fig. 5D, **Figs. S16 and S17**).

Functional annotation (Gene Ontology) of prioritized causal genes for kidney function showed enrichment for cellular metabolic processes, including organic substrate, nitrogen compound, sulfur and macromolecule metabolism (**Fig. S18A and Table S11**). The enriched pathways were consistent with the functional role of the kidney in detoxification and metabolism. Gene ontology annotation prioritized biological processes for HTN shown enrichment for angiogenesis and circulation (**Fig. S18B and Table S12**).

Taken together, kidney compartment-based eQTL(cf)s colocalization annotation of renal disease endophenotypes increased the prioritized genes by 10-fold (230 from the previous 27 (*7*)). Functional annotation highlighted the likely role of metabolism in kidney function regulation and angiogenesis in blood pressure genetics. Furthermore, eQTL(ci) enabled the discovery of driver cell types and genes previously masked by bulk tissue analysis.

### Multi-omics integrative annotation highlights important therapeutics targets for CKD and HTN

At present, the most widely used therapies for preventing kidney disease progression are inhibitors of the renin-angiotensin-aldosterone system. About 70-80% of patients with CKD take ACEi or ARBs. These drugs also effectively reduce blood pressure, and are one of the most prescribed classes of anti-hypertensive medications taken by roughly 30% of patients with HTN worldwide. Despite their remarkable efficacy and common use over the last 3 decades, the molecular mechanism of protection has not been identified as both genes are expressed by multiple cell types and genetic studies have not elucidated why these treatments are effective.

A recent SBP GWAS (*46*) identified a genetic signal on chromosome 17, rs4292, which barely passed genome wide significance (P-value_(GWAS)_=5.88E-09). Our eQTL analysis indicated that the same SNP, rs4292 was significantly associated with *ACE* expression in microdissected kidney tubule samples (**Fig. S19A**). Interestingly, we observed that the allelic dosage of C or T is markedly changed the relationship between PT fractions and *ACE* expression in human kidney proximal tubules (Fig. 6A). Meta-analysis of this eQTL association (rs4292 and *ACE*) (kidney compartments and 46 GTEx tissue (*7*), **Methods**) indicated that rs4292 had the most significant eQTL effect on *ACE* (M-value=1) in kidney, especially in the tubule fraction (Fig. 6B **and Fig. S20**). Furthermore, the signal colocalized with PT eQTL(ci) with a high confidence (Fig. 6C). An Iterative Bayesian Step-wise Selection based sum of single effect model for fine mapping analysis implemented in Susie (*53*) indicated that across the ∼2,500 SNPs located within ±1 Mb around the transcription start site of *ACE*, rs4292 has the highest posterior inclusion probability (PIP=55.5%) (**Fig. S21**) for being causal. Single cell open chromatin data indicated that rs4292 was located on a cell type specific open chromatin region only in PT around the promoter region of *ACE* (Fig. 6D). Single cell open chromatin accessibility analysis highlighted the strong correlation between open chromatin regions of *ACE* and rs4292 and other significant GWAS SNPs in PT cells (Fig. 6D **and Fig. S22A**). Finally, using gapped *k*-mer support vector machine based methods (gkm-SVM and deltaSVM) (*54, 55*), we were able to computationally quantify the likely causal regulatory effect of this variant on the transcription factor binding activity (of the 50 bp region around rs4292) significantly increased, when the C allele was replaced by allele T (deltaSVM=2.16, P-value=6.75E-07) of rs4292 (Fig. 6E). This observation is consistent with the association between rs4292 and *ACE* gene expression, for individuals with the higher dosage of allele T at the rs4292 locus, *ACE* expression levels are significantly higher in tubules (**Fig. S19A**).

**Figure 6.**
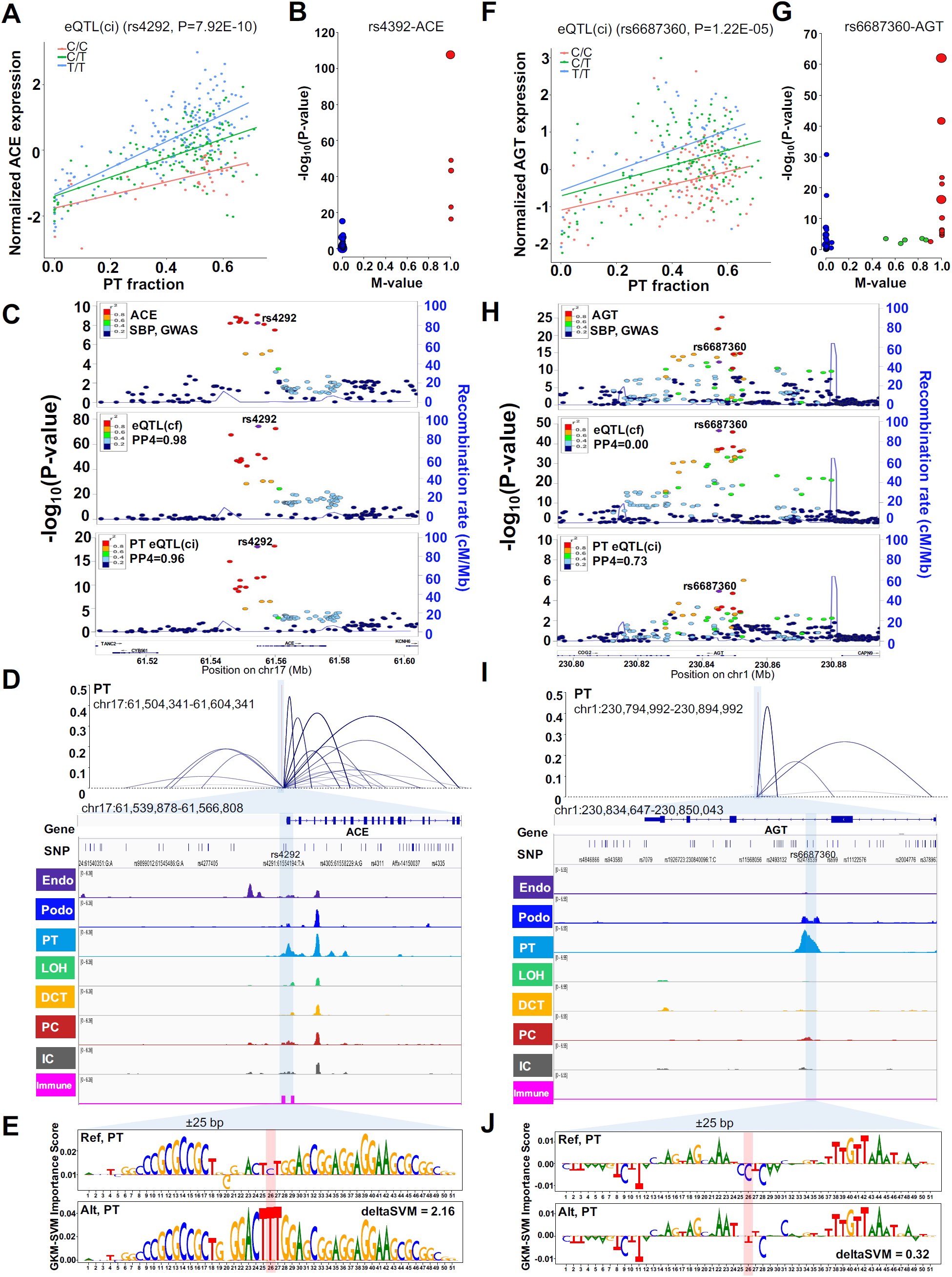
Multi-omics integrative annotation highlights important therapeutics targets for CKD and HTN. A. The y-axis is the normalized expression of *ACE* in human kidney tubules (N=356 samples), the x-axis is the PT cell fraction, each dot represents a single sample colored by their genotype C/C (red), C/T (green) and T/T (blue) at rs4292 locus. P was calculated by eQTL(ci) model. B. eQTL meta-analysis showing the association between rs4292 and *ACE* in kidney compartments and of 46 human tissues from GTEx (v7). Each dot represents one tissue, and the size represents the M-value. Red dots: M-value≥0.9, blue dots: M-value≤0.1, green dots: 0.1<M-value<0.9. The y axis shows the meta-analysis P of association in each single tissue. The x axis shows the M-value; the posterior probability of the effect in each tissue estimated by METASOFT (*61*) C. Locuszoom plots of SBP GWAS variants (top) (*46*), kidney tubule *ACE* eQTL(cf)s, and PT *ACE* eQTL(ci)s. The x-axis shows the genomic region ±100 kb around rs4292. Each data point represents a variant and the color represents the r^2^ (the degree of linkage disequilibrium). D. Top: PT cell Cicero-inferred co-accessibility between pairs of accessible chromatin sites at the *ACE* promoter. Each line is an SBP GWAS (*46*) significant SNP (P<5E-08). Single cell chromatin accessibility of human kidney single nuclear ATAC-seq data. Endo: endothelial cells, Podo: podocyte, PT: proximal tubule, LOH: loop of Henle, DCT: distal convoluted tubule, PC: collecting duct principal cells, IC: collecting duct intercalated cells, Immune: immune cells. E. Gapped *k*-mer support vector machine (GKM-SVM) method (*54*) estimated importance score for each base in the ±25 bp region surrounding rs4292 for the Ref (C) and Alt (T) alleles from the gkm-SVM model corresponding to PT cell cluster. The SNP of interest is highlighted in red color. The deltaSVM value was calculated to represent the changes of the transcription factor binding activity of this 51 bp sequence after replacing allele C to allele T of rs4292. F. The y-axis is the normalized expression of *AGT* in human kidney tubules (N=356 samples), the x-axis is the PT cell fraction, each dot represents a single sample colored by their genotype C/C (red), C/T (green) and T/T (blue) at rs6687360 locus. P was calculated by eQTL(ci) model. G. Results of eQTL from meta-analysis showing the association between rs6687360 and *AGT* using eQTLs of kidney compartments and of 46 human tissues from GTEx (v7). Each dot represents one tissue, and the size represents the M-value. Red dots: M-value≥0.9, blue dots: M-value≤0.1, green dots: 0.1<M-value<0.9. The y-axis shows the meta-analysis P of association in each single tissue. The x-axis shows the M-value; the posterior probability of the effect in each tissue estimated by METASOFT (*61*). H. Locuszoom plots of SBP GWAS variants (*46*), kidney tubule eQTL(cf)s, and PT *AGT* eQTL(ci)s. The x-axis shows the genomic region ±100 kb around rs6687360. Each data point represents a variant and the color represent the r^2^ (the degree of LD). I. Top: PT cell Cicero-inferred co-accessibility between pairs of accessible chromatin sites at the AGT gene body region. Each line is an SBP GWAS (46) significant SNP (P<5E-08). Single cell chromatin accessibility of human kidney single nuclear ATAC-seq data. Endo: endothelial cells, Podo: podocyte, PT: proximal tubule, LOH: loop of Henle, DCT: distal convoluted tubule, PC: collecting duct principal cells, IC: collecting duct intercalated cells, Immune: immune cells. The showing genomic region is chr1:230,834,647-230,850,043. J. Gapped *k*-mer support vector machine (GKM-SVM) method (*54*) estimated importance score for each base in the ±25 bp region surrounding rs6687360 for the Ref and Alt alleles from the gkm-SVM model corresponding to PT cell cluster. The SNP of interest is highlighted in red color. The deltaSVM value was calculated to represent the changes of the transcription factor binding activity of this 51 bp sequence after replacing allele C to allele T of rs6687360.

To our surprise, we found that our analysis identified not only *ACE*, but also *AGT*, encoding the primary substrate for production of angiotensin II. The SBP GWAS (*46*) have identified a significant signal at rs6687360 (P_GWAS(rs668(7360)_=4.72E-13). Kidney tubule eQTL data highlighted a strong association with *AGT* expression (P_eQTL_=4.83E-44) (**Fig. S19B**). Allelic dosage of C at the rs6687360 variant, significantly reduced the correlation between PT fraction and *AGT* expression with P-value=1.22E-05 (Fig. 6F). Interestingly, although *AGT* is expressed in multiple tissues and cell types (*34*), meta-analysis of the eQTL rs6687360-*AGT* in kidney compartments studies (*7*) and 46 GTEx tissues (**Methods**) indicated that rs6687360 was most significantly associated with *AGT* (M-value=1) in the kidney, especially in the tubule compartment (Fig. 6G **and Fig. S23**). We did not observe colocalization with the genetic signal using bulk eQTL data from tubule (PP4=0.00), but found a strong colocalization signal between PT eQTL(ci) and the genetic signal, highlighting the shared genetic casual variants between the GWAS and PT eQTL(ci) (PP4=0.73) (Fig. 6H). We only observed a single open chromatin region of this area in the intronic region of *AGT* in PT cells (Fig. 6I). Single-cell open chromatin co-accessibility analysis showed an interaction between opening regions of *AGT* and rs6687360 (and other significant GWAS SNPs in PT cells (Fig. 6I **and Fig. S22B**)). In addition, the deltaSVM analysis indicated the TF binding activity of ±25 bp region around rs6687360 increased as allele C changed to T on rs6687360 in PT cells (Fig. 6J). The observation was consistent with tubule eQTL effect (**Fig. S19B**), samples with the higher dosage of T at rs6687360 locus had higher *AGT* expression.

The discovery of more than 200 prioritized genes for kidney function and hypertension allowed us to identify targets for potential repurposing of drugs. Using the Drug Gene Interaction Database (DGIdb v3.0, https://www.dgidb.org), we identified 54 genes that can be targeted by already FDA-approved drugs (**Table S15**). In addition, to the large number of drugs that targeted *ACE* and *AGT* with clinically approval drugs, several drugs directly target prioritized genes, such as cilastatin that inhibits the kidney dipeptidase (*DPEP1*) (**Table S15**). *DPEP1* was nominated as a kidney disease gene based on our eQTL annotation. The *DPEP1* inhibitor cilastatin showed protection in preliminary studies from acute toxic kidney injury in animal models (*56*). Caspase9 has also been prioritized as a CKD gene, with already available inhibitors, such as emricasan and nivocasan (**Table S15**). Several genes from Solute Carrier Family (SLC) have been identified in our study that might be particularly amenable for pharmacological targeting, such as *SLC2A4* (GLUT4), *SLC47A1* and *SLC15A2*(*57*), in addition to the key regulator of the Notch signaling; *APH1A*, hedgehog signaling; *SMO* and *TGFB1* (**Table S15 and S16**).

Overall, our combined eQTL(ci) and snATAC-based GWAS analysis had defined the genetic and mechanistic basis for ACEi and ARBs; the most commonly used anti-hypertensive and kidney protective drugs, and prioritized a large number of mostly approved compounds that can be repurposed for the treatment of kidney disease and hypertension.

## Discussion

Here, we generated a comprehensive multi-omics dataset and used orthogonal analytical approaches to annotate renal endophenotypes. Our analysis prioritized more than 182 likely causal genes for kidney function and 88 for hypertension, potentially defining the core genes for these conditions in the kidney. We showed the critical cell type convergence of endophenotypes, such as kidney function and metabolite heritability enrichment in PT and SBP heritability enrichment in endothelial cells and distal tubule segments. Functional annotation indicated the likely role of metabolism in kidney disease and angiogenesis in hypertension. Finally, our analysis clarified the key genetic underpinning of action for the most commonly used antihypertensive and renal protective drugs, as well as highlighted a large number of additional drugs that could potentially be effective for HTN and CKD.

While GWAS studies have been extremely powerful in disease associated loci discovery, signals at these sites have rarely been translated into molecular mechanisms and therapeutics. One of the most important computational methods used to annotate GWAS signals is eQTL analysis. Limited availability of kidney eQTL datasets has been an important limitation of renal related traits GWAS annotation. With the increased sample size provided in this study, we were able to double the number of identified eQTLs.

Another key limitation of the eQTL studies has been the heterogenous cell types in the bulk tissue samples. We found that while PEER factors have been able to account for cell fraction heterogeneity, additional adjustment using computational deconvolution could further improve the model. We took this a step further and identified eQTL(ci) that define genotype-cell interaction (cell-type) effects. We cataloged SNP-gene pairs by examining the interaction of cell fraction and genetic variations in a large number of kidney compartment samples. We also noted some limitations of the eQTL(ci) approach. First, the approach is limited by the quality of the single cell gene expression data. While human kidney single cell RNA-seq data are now available, we found that the higher quality mice dataset generated more reliable information on cell fractions (*16, 58*). Second, the co-linearity of cell fractions in tissue samples, such as the loss of kidney parenchyma, is associated with a relative increase in proportion of mesenchymal and immune cells, and so these methods are unable to distinguish between such closely related cell fractions. Finally, the analysis appeared to be more robust for the abundant cell types of each tissue. While eQTL(ci) is a new and exciting method, the model has not been experimentally validated. Here, we used snATAC-seq data to understand whether eQTL(ci) could be linked to cell-type specific open chromatin regions. Our data supports the results from eQTL(ci)s as they showed enrichment for cell type specific regulatory regions.

Our combined analyses that used LDSC (for scRNA-seq and snATAC-seq data), MAGMA and eQTL(ci) coloc have provided several meaningful insights into driver cell types for kidney disease-associated endophenotypes. We demonstrated that eGFR heritability is primarily derived from eQTL influence in PT cells where metabolite levels, such as serum uric acid, are also regulated by PT functions. In contrast, SBP was associated with genes expressed specifically in endothelial cells and DCT cells. The cell type enrichment of endophenotypes appears to be robust and highly consistent using different methods and datasets.

Finally, our results provide insights into previously cryptic disease mechanisms and therapeutic targeting opportunities. Inhibitors of the renin-angiotensin-aldosterone system, such as the ACEi and ARBs, are the most commonly used medication class for SBP and preventing kidney disease progression. Yet, we lacked human genetic data supporting the therapeutic benefits of ACEi and ARB in CKD or hypertension. The *ACE* locus passed genome wide significance in a large SBP GWAS and our kidney eQTL annotation provided critical information for the annotation of this locus and highlighted its association with *ACE* expression in PTs. Our orthogonal analyses including snATAC and deltaSVM nominated rs4292 for this effect.

It is also important to note that GWAS variants at the *AGT* locus were robustly associated with SBP, but the associations of genetic variants with gene expression using bulk eQTL datasets were not sufficient to prioritize the causal gene for this locus. It was really the interaction effect between genetic variants and PT cell fraction variations identified from PT eQTL(ci) and snATAC data that had helped with the target gene and cell type prioritization. Mouse model experiments are consistent with our results, notably, a linear correlation between *AGT* levels and blood pressure has been previously established in mouse models (*59*). *AGT* made in PT is shed into the tubular fluid (*60*), where it co-localizes with *ACE*, which is highly expressed on the brush border on the tubular surface of proximal epithelia. Overall, our study highlights the critical role of the kidney in blood pressure regulation and raises the possibility that these variants might be useful for future precision therapeutic approaches, such as to understand differences in response to ACEi and ARB.

In summary, we present an integrated multi-omics study for functional interpretation of common variants associated with renal traits using kidney compartment-based eQTL and single-cell resolution open chromatin identification. The application of multiple orthogonal computational methods was critical to prioritize more than 200 genes for eGFR and SBP, highlighted the key disease-associated variants, cell types and disease pathomechanism, clarified mechanism of widely used therapeutics and highlighted new potential drugs for these kidney related diseases.

## Author contributions

KS and XS conceived, planned and oversaw the study, and wrote the manuscript. ZMa and JW conducted the human kidney snATAC-seq experiment. XS analyzed data with the help of HL, CQ, ZMi. MJS, MP, MKS, KD, SSP, TLE, JNH, AMH, ML, BV, TC, CDB, KS assisted with data generation and manuscript revision.

## Data and materials availability

The RNA-seq, eQTL data and human kidney snATAC-seq data is publicly available at the diabetes epigenome atlas www.diabetesepigenome.org. No consent was obtained to share individual-level genotype data.

## Supplementary Materials

Materials and Methods

Figs. S1 to S28

Tables S1 to S16

References (1-64)

