## Supplementary Figures for "Mapping the genetic architecture of human traits to cell types in the kidney identifies mechanisms of disease and potential treatments"

**This PDF file include:**

Figs. S1 to S28
References (1-64)

**Other Supplementary Materials for this manuscript include the following:**

Tables S1 to S16 (Excel)

**Supplementary figure 1**

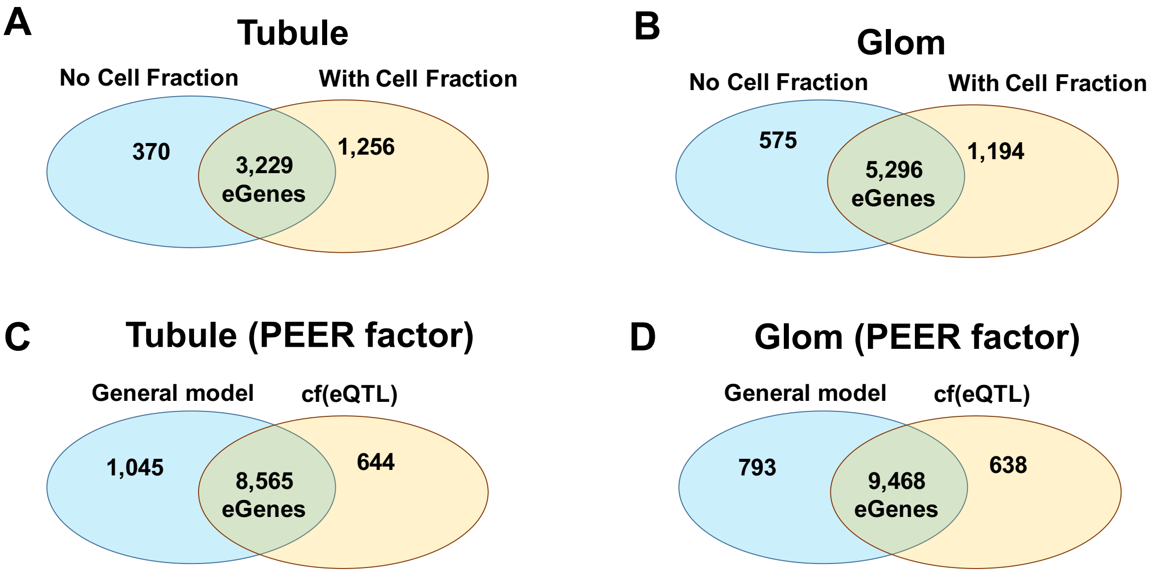


**Fig. S1. The comparison of eGenes identified by different models** A). Venn diagram illustrating the overlap of eGenes identified by the general eQTL model (with PEER factors were excluded) and the eQTL(cf) model (without PEER factors) in 356 human tubule samples. B). Venn diagram illustrating the overlap of eGenes identified by the general eQTL model (without PEER factors) and the eQTL(cf) model (without PEER factors) in 303 human glomerular samples. C). Venn diagram illustrating the overlap of eGenes identified by the general eQTL model and the eQTL(cf) model in 356 human tubule samples. D). Venn diagram illustrating the overlap of eGenes identified by the general eQTL model and the eQTL(cf) model in 303 human glomerular samples.

**Supplementary figure 2**

**A**

**Tubuli**

| | PT | ALOH | DCT | B-IC | CNT | DC | Granul | Treg | NK | Th17 | CD8+ T | CD4+ T | Macro | Fibrosis | Batch | Site | Paired-end | Age | Gender | RIN | PC1 | PC2 | PC3 | PC4 | PC5 |
|---|---|---|---|---|---|---|---|---|---|---|---|---|---|---|---|---|---|---|---|---|---|---|---|---|---|
| PEER1 | 0.37 | -0.14 | -0.17 | -0.07 | -0.29 | -0.43 | 0.07 | -0.02 | -0.20 | -0.21 | -0.14 | -0.23 | -0.17 | -0.18 | -0.15 | -0.15 | -0.19 | -0.05 | 0.06 | -0.31 | 0.13 | 0.14 | 0.03 | 0.13 | -0.03 |
| PEER2 | 0.09 | -0.01 | 0.00 | -0.18 | 0.09 | -0.02 | -0.15 | -0.14 | -0.12 | -0.11 | -0.16 | -0.02 | -0.10 | -0.18 | 0.67 | 0.49 | 0.08 | 0.03 | -0.16 | 0.06 | -0.36 | -0.42 | -0.24 | -0.31 | 0.19 |
| PEER3 | 0.43 | 0.01 | -0.26 | -0.56 | 0.06 | -0.27 | -0.17 | -0.24 | -0.38 | -0.56 | -0.52 | -0.03 | -0.36 | -0.30 | -0.09 | 0.10 | 0.10 | 0.14 | 0.02 | 0.21 | -0.09 | -0.06 | -0.06 | 0.03 | -0.04 |
| PEER4 | 0.48 | -0.03 | -0.08 | -0.30 | -0.24 | -0.50 | -0.24 | -0.24 | -0.37 | -0.40 | -0.21 | -0.45 | -0.36 | -0.43 | 0.58 | 0.24 | 0.33 | 0.03 | -0.03 | 0.12 | -0.21 | -0.25 | -0.14 | -0.19 | 0.09 |
| PEER5 | 0.48 | 0.40 | 0.26 | -0.48 | -0.63 | -0.56 | -0.30 | -0.29 | -0.51 | -0.22 | -0.44 | -0.43 | -0.52 | -0.44 | 0.10 | 0.05 | -0.03 | -0.15 | 0.13 | -0.03 | -0.06 | -0.07 | -0.11 | 0.00 | 0.01 |
| PEER6 | 0.74 | 0.08 | -0.09 | -0.64 | -0.44 | -0.74 | -0.34 | -0.39 | -0.66 | -0.56 | -0.56 | -0.46 | -0.65 | -0.67 | 0.15 | 0.09 | -0.18 | -0.11 | -0.06 | -0.05 | -0.14 | -0.13 | -0.10 | -0.05 | 0.06 |
| PEER7 | -0.69 | -0.18 | -0.10 | 0.57 | 0.59 | 0.71 | 0.30 | 0.42 | 0.70 | 0.38 | 0.61 | 0.35 | 0.63 | 0.55 | -0.23 | -0.13 | -0.14 | 0.16 | 0.03 | -0.02 | 0.14 | 0.13 | 0.11 | 0.03 | -0.04 |
| PEER8 | -0.27 | 0.05 | 0.14 | 0.10 | 0.06 | 0.14 | 0.22 | 0.12 | 0.17 | 0.22 | 0.02 | 0.20 | 0.16 | 0.18 | -0.18 | 0.10 | 0.10 | 0.08 | -0.04 | -0.11 | -0.10 | -0.13 | -0.10 | -0.06 | -0.04 |
| PEER9 | -0.12 | -0.20 | 0.02 | 0.20 | 0.05 | 0.12 | 0.08 | 0.15 | 0.20 | 0.12 | 0.35 | 0.02 | 0.18 | 0.00 | 0.03 | 0.00 | -0.18 | 0.00 | 0.03 | 0.09 | -0.17 | -0.14 | -0.06 | -0.05 | 0.16 |
| PEER10 | 0.34 | 0.37 | -0.02 | -0.24 | -0.45 | -0.23 | -0.18 | -0.11 | -0.24 | -0.16 | -0.15 | -0.01 | -0.23 | -0.22 | 0.05 | -0.04 | -0.07 | -0.13 | -0.10 | 0.11 | -0.08 | -0.09 | -0.07 | -0.03 | 0.04 |
| PEER11 | -0.32 | 0.40 | 0.39 | 0.18 | -0.13 | 0.08 | 0.16 | -0.02 | 0.14 | 0.31 | 0.13 | 0.01 | 0.13 | 0.20 | 0.14 | 0.06 | 0.23 | 0.14 | -0.05 | 0.12 | 0.04 | 0.00 | 0.04 | -0.07 | -0.09 |
| PEER12 | 0.49 | -0.59 | -0.37 | -0.13 | 0.09 | -0.25 | -0.12 | 0.03 | -0.07 | -0.48 | -0.03 | -0.22 | -0.08 | -0.15 | 0.18 | 0.11 | 0.05 | 0.07 | -0.01 | 0.05 | -0.12 | -0.11 | -0.06 | -0.04 | 0.08 |
| PEER13 | -0.37 | 0.24 | 0.57 | 0.16 | -0.12 | 0.10 | 0.26 | -0.03 | 0.07 | 0.53 | 0.07 | 0.12 | 0.12 | 0.03 | 0.07 | 0.07 | 0.05 | -0.11 | 0.07 | 0.14 | 0.01 | -0.02 | 0.00 | -0.07 | 0.05 |
| PEER14 | 0.31 | -0.24 | -0.39 | -0.08 | 0.01 | -0.12 | -0.02 | 0.02 | -0.10 | -0.29 | -0.13 | 0.04 | -0.10 | 0.09 | -0.04 | 0.04 | -0.02 | -0.02 | 0.13 | -0.07 | 0.13 | 0.10 | 0.04 | 0.03 | 0.04 |
| PEER15 | -0.49 | -0.38 | 0.35 | 0.49 | 0.17 | 0.50 | 0.27 | 0.39 | 0.55 | 0.51 | 0.42 | 0.20 | 0.49 | 0.44 | -0.05 | -0.01 | 0.20 | -0.07 | -0.06 | 0.03 | 0.14 | 0.10 | 0.03 | 0.02 | -0.07 |
| PEER16 | 0.39 | 0.08 | -0.15 | -0.37 | -0.21 | -0.38 | -0.07 | -0.27 | -0.37 | -0.31 | -0.42 | -0.12 | -0.28 | -0.26 | 0.13 | 0.05 | 0.08 | 0.21 | -0.09 | -0.19 | 0.06 | 0.04 | 0.10 | -0.06 | -0.06 |
| PEER17 | -0.28 | -0.21 | -0.09 | 0.34 | 0.37 | 0.33 | 0.23 | 0.09 | 0.19 | 0.25 | 0.25 | 0.23 | 0.29 | 0.43 | -0.16 | -0.24 | -0.14 | -0.07 | 0.03 | -0.09 | 0.15 | 0.14 | 0.09 | 0.05 | -0.04 |
| PEER18 | 0.44 | -0.24 | -0.41 | -0.23 | 0.07 | -0.27 | -0.17 | -0.27 | -0.19 | -0.51 | -0.18 | -0.12 | -0.20 | -0.20 | 0.09 | 0.16 | 0.15 | -0.06 | 0.12 | -0.15 | 0.04 | 0.05 | 0.06 | 0.01 | 0.03 |
| PEER19 | 0.07 | 0.03 | -0.17 | -0.14 | -0.14 | -0.05 | 0.00 | 0.03 | -0.09 | -0.14 | -0.23 | 0.06 | -0.12 | -0.17 | -0.26 | -0.17 | -0.33 | 0.00 | -0.15 | -0.01 | -0.02 | 0.06 | 0.12 | 0.08 | 0.01 |
| PEER20 | 0.30 | -0.31 | -0.34 | -0.12 | 0.15 | -0.14 | -0.09 | -0.15 | -0.17 | -0.33 | -0.18 | -0.15 | -0.16 | -0.05 | 0.04 | 0.22 | 0.29 | -0.03 | -0.15 | -0.11 | -0.22 | -0.18 | -0.05 | -0.04 | -0.08 |
| PEER21 | -0.06 | 0.16 | -0.04 | 0.09 | -0.04 | -0.04 | -0.07 | 0.08 | 0.07 | -0.11 | 0.10 | -0.15 | -0.02 | -0.03 | 0.12 | 0.07 | 0.11 | -0.04 | -0.14 | 0.07 | -0.15 | -0.17 | -0.08 | -0.14 | 0.10 |
| PEER22 | -0.22 | 0.31 | 0.40 | 0.19 | -0.09 | 0.10 | 0.19 | 0.15 | 0.07 | 0.26 | 0.13 | 0.16 | 0.13 | 0.07 | 0.22 | 0.11 | 0.39 | -0.20 | -0.11 | 0.13 | 0.02 | 0.03 | 0.05 | 0.02 | -0.15 |
| PEER23 | -0.07 | -0.11 | 0.03 | 0.04 | 0.24 | 0.03 | -0.02 | -0.06 | -0.03 | 0.18 | 0.08 | -0.07 | -0.03 | 0.08 | -0.17 | -0.01 | -0.21 | -0.04 | 0.15 | -0.07 | -0.01 | -0.04 | -0.06 | -0.03 | 0.01 |
| PEER24 | 0.01 | -0.25 | -0.06 | 0.16 | 0.15 | 0.20 | 0.11 | 0.19 | 0.21 | 0.00 | 0.18 | 0.14 | 0.13 | 0.12 | -0.09 | -0.18 | -0.37 | -0.03 | 0.04 | -0.02 | 0.22 | 0.19 | 0.10 | 0.04 | 0.15 |
| PEER25 | 0.39 | 0.09 | -0.20 | -0.32 | -0.29 | -0.23 | -0.17 | 0.01 | -0.20 | -0.42 | -0.29 | -0.09 | -0.23 | -0.25 | 0.26 | 0.07 | 0.29 | 0.05 | -0.06 | -0.11 | 0.16 | 0.12 | 0.05 | 0.02 | -0.09 |
| PEER26 | 0.35 | -0.12 | -0.39 | -0.20 | 0.00 | -0.24 | -0.28 | -0.14 | -0.12 | -0.47 | -0.12 | -0.11 | -0.11 | -0.17 | -0.05 | 0.08 | -0.10 | 0.12 | 0.05 | -0.17 | -0.11 | -0.18 | -0.18 | -0.13 | 0.04 |
| PEER27 | -0.31 | -0.08 | 0.12 | 0.43 | 0.09 | 0.32 | 0.19 | 0.27 | 0.44 | 0.25 | 0.36 | 0.16 | 0.34 | 0.30 | 0.04 | -0.01 | 0.19 | -0.05 | -0.11 | -0.02 | 0.05 | 0.04 | 0.05 | 0.01 | -0.10 |
| PEER28 | 0.02 | -0.07 | 0.06 | -0.05 | -0.02 | -0.16 | -0.06 | -0.30 | -0.14 | 0.15 | -0.05 | -0.09 | -0.13 | 0.06 | -0.04 | -0.05 | -0.15 | 0.04 | -0.22 | -0.03 | 0.27 | 0.25 | 0.14 | 0.07 | 0.03 |
| PEER29 | -0.08 | 0.35 | 0.24 | -0.05 | -0.15 | -0.02 | 0.01 | -0.02 | -0.10 | 0.19 | 0.00 | -0.11 | -0.04 | -0.07 | 0.02 | 0.00 | 0.21 | -0.03 | 0.21 | 0.05 | -0.21 | -0.19 | -0.05 | -0.11 | -0.01 |
| PEER30 | 0.11 | -0.23 | -0.05 | 0.03 | -0.10 | 0.04 | -0.01 | -0.03 | 0.10 | -0.07 | -0.02 | 0.07 | 0.05 | -0.04 | 0.04 | 0.05 | 0.28 | 0.04 | -0.18 | -0.04 | -0.11 | -0.06 | 0.00 | 0.04 | -0.08 |
| PEER31 | -0.56 | 0.20 | 0.34 | 0.44 | 0.22 | 0.40 | 0.23 | 0.11 | 0.33 | 0.55 | 0.41 | 0.18 | 0.31 | 0.36 | -0.04 | -0.22 | -0.08 | -0.15 | 0.15 | 0.13 | 0.25 | 0.23 | 0.11 | 0.08 | -0.02 |
| PEER32 | 0.47 | 0.18 | -0.11 | -0.46 | -0.29 | -0.40 | -0.31 | -0.32 | -0.50 | -0.28 | -0.43 | -0.14 | -0.53 | -0.39 | 0.22 | 0.17 | 0.16 | 0.02 | 0.11 | -0.10 | -0.16 | -0.16 | -0.16 | -0.04 | -0.02 |
| PEER33 | -0.21 | 0.08 | 0.03 | 0.06 | 0.13 | 0.12 | 0.12 | 0.04 | -0.01 | 0.17 | 0.06 | 0.10 | 0.03 | 0.11 | -0.13 | 0.01 | -0.01 | 0.01 | -0.11 | 0.06 | -0.01 | 0.02 | 0.06 | 0.03 | -0.07 |
| PEER34 | 0.30 | -0.32 | -0.27 | -0.06 | 0.06 | -0.04 | -0.07 | -0.03 | 0.02 | -0.34 | -0.03 | -0.04 | -0.06 | -0.05 | 0.17 | 0.19 | 0.60 | -0.04 | 0.04 | -0.11 | 0.03 | 0.04 | 0.06 | 0.01 | -0.12 |
| PEER35 | -0.05 | 0.20 | 0.14 | -0.03 | -0.15 | 0.12 | -0.17 | -0.09 | 0.06 | 0.08 | 0.04 | -0.05 | -0.05 | -0.06 | 0.18 | 0.26 | 0.57 | -0.05 | -0.18 | -0.04 | -0.17 | -0.12 | -0.03 | -0.03 | -0.14 |

**B**

**Glomeruli**

| | GEC | Endo | Podo | Granul | Treg | NK | Th17 | CD8+ T | CD4+ T | Macro | Sclerosis | Batch | Paired-end | Age | Gender | RIN | Site | PC1 | PC2 | PC3 | PC4 | PC5 |
|---|---|---|---|---|---|---|---|---|---|---|---|---|---|---|---|---|---|---|---|---|---|---|
| PEER1 | 0.45 | -0.31 | 0.22 | -0.30 | -0.03 | 0.14 | -0.01 | -0.06 | 0.04 | -0.24 | -0.03 | 0.28 | -0.16 | 0.04 | -0.10 | 0.43 | 0.48 | -0.35 | -0.42 | -0.24 | -0.32 | 0.27 |
| PEER2 | 0.23 | -0.10 | 0.16 | -0.16 | 0.04 | 0.30 | 0.04 | -0.11 | 0.08 | 0.06 | 0.18 | -0.35 | -0.38 | 0.01 | -0.06 | 0.24 | -0.05 | 0.02 | 0.03 | 0.02 | 0.01 | 0.08 |
| PEER3 | 0.58 | -0.34 | 0.17 | -0.21 | 0.19 | 0.32 | -0.25 | -0.23 | 0.21 | -0.13 | 0.11 | -0.45 | -0.11 | 0.21 | -0.19 | 0.04 | 0.29 | -0.14 | -0.13 | -0.06 | -0.05 | 0.04 |
| PEER4 | 0.02 | 0.01 | 0.18 | 0.18 | -0.27 | -0.59 | -0.28 | 0.29 | -0.05 | -0.19 | -0.24 | -0.20 | -0.24 | -0.10 | 0.02 | 0.27 | -0.03 | -0.03 | 0.01 | 0.06 | 0.02 | -0.01 |
| PEER5 | -0.04 | 0.15 | -0.23 | -0.03 | 0.27 | -0.04 | -0.05 | 0.11 | 0.04 | 0.11 | -0.13 | -0.17 | -0.06 | -0.07 | -0.02 | -0.31 | 0.13 | -0.08 | -0.09 | -0.09 | -0.04 | -0.02 |
| PEER6 | -0.55 | 0.61 | -0.62 | 0.38 | 0.16 | -0.20 | -0.06 | 0.02 | 0.09 | 0.72 | 0.23 | 0.01 | 0.10 | -0.06 | 0.16 | -0.21 | -0.34 | 0.14 | 0.19 | 0.18 | 0.12 | -0.07 |
| PEER7 | -0.25 | 0.26 | -0.42 | 0.22 | 0.10 | 0.11 | 0.10 | -0.07 | -0.11 | 0.53 | 0.29 | -0.31 | -0.02 | 0.15 | 0.06 | -0.36 | -0.20 | 0.39 | 0.40 | 0.25 | 0.19 | -0.08 |
| PEER8 | -0.06 | 0.08 | -0.05 | -0.22 | 0.16 | 0.29 | 0.19 | -0.11 | 0.06 | 0.12 | 0.16 | 0.18 | 0.10 | 0.07 | 0.14 | -0.02 | 0.01 | 0.08 | 0.05 | -0.05 | 0.02 | 0.02 |
| PEER9 | -0.63 | 0.56 | -0.33 | 0.44 | -0.12 | -0.62 | -0.04 | 0.41 | -0.09 | 0.38 | -0.07 | 0.07 | 0.13 | -0.11 | 0.11 | -0.09 | -0.31 | 0.09 | 0.13 | 0.13 | 0.09 | -0.04 |
| PEER10 | -0.11 | 0.12 | 0.09 | 0.11 | -0.14 | -0.16 | 0.01 | 0.01 | -0.02 | -0.07 | -0.04 | -0.03 | -0.07 | -0.05 | -0.05 | -0.07 | 0.07 | -0.06 | -0.07 | -0.01 | -0.06 | -0.01 |
| PEER11 | 0.41 | -0.17 | -0.05 | 0.02 | -0.07 | 0.01 | -0.23 | -0.08 | 0.10 | -0.12 | -0.08 | -0.27 | -0.13 | 0.12 | 0.03 | -0.20 | 0.21 | -0.16 | -0.17 | -0.11 | -0.05 | -0.09 |
| PEER12 | 0.32 | -0.08 | -0.16 | -0.17 | 0.00 | 0.10 | -0.03 | 0.03 | -0.13 | -0.10 | 0.08 | -0.12 | -0.16 | -0.09 | 0.06 | 0.04 | 0.00 | 0.11 | 0.10 | 0.00 | 0.11 | -0.04 |
| PEER13 | 0.03 | 0.16 | -0.33 | 0.24 | -0.08 | -0.21 | -0.27 | 0.02 | 0.17 | 0.14 | -0.08 | -0.21 | -0.20 | -0.13 | 0.24 | -0.04 | -0.03 | 0.16 | 0.15 | 0.04 | 0.10 | 0.02 |
| PEER14 | -0.04 | 0.05 | -0.26 | 0.32 | 0.01 | -0.09 | -0.23 | -0.15 | 0.27 | 0.10 | -0.08 | 0.04 | 0.00 | -0.07 | 0.02 | 0.03 | 0.07 | 0.10 | 0.10 | 0.01 | 0.07 | -0.03 |
| PEER15 | -0.19 | 0.11 | 0.10 | 0.15 | -0.04 | -0.14 | -0.07 | 0.09 | 0.06 | 0.15 | 0.00 | -0.10 | -0.34 | -0.03 | 0.00 | 0.03 | -0.04 | 0.04 | 0.00 | -0.08 | 0.00 | 0.03 |
| PEER16 | -0.14 | -0.01 | 0.06 | -0.08 | 0.06 | 0.08 | 0.13 | -0.09 | -0.01 | 0.04 | 0.13 | 0.21 | 0.23 | 0.05 | 0.12 | 0.07 | 0.02 | 0.08 | 0.06 | -0.06 | 0.05 | 0.01 |
| PEER17 | 0.56 | -0.39 | 0.41 | -0.26 | -0.15 | 0.13 | -0.12 | -0.02 | -0.02 | -0.53 | -0.15 | -0.46 | -0.50 | 0.13 | -0.07 | 0.08 | 0.12 | -0.21 | -0.18 | -0.10 | -0.01 | -0.03 |
| PEER18 | 0.06 | 0.13 | -0.46 | -0.23 | 0.33 | 0.06 | 0.27 | 0.08 | -0.20 | 0.15 | 0.09 | 0.13 | 0.43 | -0.04 | 0.13 | -0.08 | -0.04 | 0.00 | -0.03 | 0.05 | -0.09 | -0.05 |
| PEER19 | -0.44 | 0.15 | 0.05 | 0.14 | -0.08 | 0.04 | 0.22 | -0.11 | 0.12 | 0.10 | 0.09 | 0.25 | 0.16 | -0.07 | -0.15 | -0.01 | -0.11 | 0.13 | 0.11 | 0.05 | 0.02 | -0.02 |
| PEER20 | -0.49 | 0.34 | -0.26 | 0.22 | 0.16 | -0.14 | 0.02 | 0.09 | -0.01 | 0.49 | 0.02 | 0.26 | 0.19 | -0.14 | -0.01 | 0.01 | -0.17 | -0.03 | -0.05 | -0.07 | -0.04 | -0.04 |
| PEER21 | 0.06 | 0.05 | -0.13 | 0.16 | 0.05 | -0.04 | -0.08 | 0.04 | 0.00 | 0.01 | -0.12 | -0.01 | 0.08 | -0.14 | -0.28 | -0.12 | 0.01 | -0.04 | 0.01 | 0.06 | 0.04 | -0.03 |
| PEER22 | 0.09 | 0.00 | -0.04 | 0.05 | 0.03 | -0.12 | -0.13 | 0.04 | 0.08 | -0.06 | -0.12 | -0.02 | 0.04 | -0.23 | 0.14 | -0.07 | -0.04 | 0.11 | 0.16 | 0.10 | 0.16 | -0.11 |
| PEER23 | 0.06 | -0.17 | 0.26 | -0.01 | -0.20 | 0.06 | -0.08 | 0.11 | -0.09 | -0.04 | 0.08 | -0.24 | -0.17 | 0.10 | 0.27 | -0.12 | 0.00 | -0.19 | -0.20 | -0.16 | -0.07 | 0.02 |
| PEER24 | 0.01 | 0.03 | -0.02 | -0.02 | 0.05 | 0.02 | -0.13 | 0.07 | 0.01 | 0.16 | 0.07 | -0.11 | -0.22 | 0.07 | 0.29 | -0.01 | 0.00 | -0.20 | -0.20 | -0.09 | -0.10 | -0.03 |
| PEER25 | -0.10 | 0.00 | 0.24 | 0.06 | -0.23 | -0.05 | -0.11 | 0.03 | 0.06 | -0.03 | 0.08 | -0.19 | -0.31 | 0.10 | 0.22 | 0.00 | -0.04 | -0.22 | -0.21 | -0.13 | -0.08 | 0.00 |
| PEER26 | 0.41 | -0.35 | 0.39 | -0.07 | -0.24 | -0.01 | -0.31 | 0.02 | 0.09 | -0.37 | -0.09 | -0.33 | -0.37 | -0.10 | -0.02 | 0.08 | 0.18 | -0.16 | -0.12 | -0.11 | 0.02 | 0.10 |
| PEER27 | -0.01 | -0.02 | 0.14 | -0.02 | 0.04 | 0.11 | 0.03 | -0.11 | 0.00 | 0.00 | 0.03 | -0.07 | -0.09 | 0.07 | -0.26 | 0.07 | -0.06 | 0.04 | 0.05 | -0.05 | 0.10 | 0.01 |
| PEER28 | 0.11 | -0.05 | 0.12 | 0.04 | -0.14 | -0.04 | -0.01 | -0.12 | 0.10 | -0.26 | -0.09 | -0.16 | -0.30 | -0.03 | 0.24 | 0.05 | 0.00 | -0.21 | -0.13 | -0.03 | 0.03 | 0.03 |
| PEER29 | 0.10 | 0.04 | -0.05 | 0.04 | 0.08 | 0.02 | -0.08 | 0.01 | 0.02 | -0.10 | -0.09 | -0.04 | 0.09 | -0.31 | -0.04 | -0.06 | 0.04 | -0.08 | 0.01 | 0.02 | 0.15 | -0.04 |
| PEER30 | -0.12 | 0.08 | -0.10 | 0.10 | 0.07 | -0.13 | 0.09 | -0.04 | -0.03 | 0.18 | 0.13 | 0.11 | 0.00 | 0.09 | -0.16 | 0.11 | -0.05 | 0.07 | 0.17 | 0.28 | 0.09 | 0.02 |
| PEER31 | -0.07 | 0.07 | -0.15 | 0.03 | 0.04 | 0.00 | 0.04 | 0.04 | -0.06 | 0.06 | 0.11 | 0.11 | 0.30 | 0.22 | -0.03 | -0.19 | -0.05 | 0.32 | 0.25 | 0.12 | 0.02 | -0.03 |
| PEER32 | 0.06 | 0.04 | 0.07 | 0.02 | -0.13 | -0.10 | -0.01 | -0.04 | 0.05 | -0.16 | -0.13 | 0.02 | -0.05 | -0.15 | -0.01 | 0.11 | 0.01 | -0.04 | 0.04 | 0.07 | 0.09 | 0.02 |
| PEER33 | -0.07 | 0.16 | -0.21 | 0.12 | 0.20 | -0.06 | -0.01 | -0.01 | 0.03 | 0.13 | -0.01 | 0.19 | 0.34 | -0.17 | -0.01 | 0.03 | -0.05 | 0.29 | 0.28 | 0.24 | 0.08 | -0.05 |
| PEER34 | 0.25 | -0.18 | 0.13 | 0.04 | -0.07 | -0.04 | -0.16 | -0.02 | 0.06 | -0.23 | -0.10 | -0.11 | 0.08 | -0.20 | -0.25 | 0.11 | 0.16 | -0.29 | -0.20 | -0.05 | 0.02 | 0.12 |
| PEER35 | -0.08 | 0.01 | 0.07 | 0.02 | -0.18 | 0.07 | 0.03 | -0.09 | 0.07 | -0.20 | -0.08 | -0.10 | -0.27 | -0.09 | -0.25 | -0.06 | -0.01 | 0.13 | 0.12 | 0.07 | 0.05 | 0.02 |

**Fig. S2. The Pearson's correlation between known clinical, technical variables, cell fractions and 35 PEER factors estimated from RNA-seq data of** A). 356 tubule samples B). 303 glomerular samples.

**Supplementary figure 3**

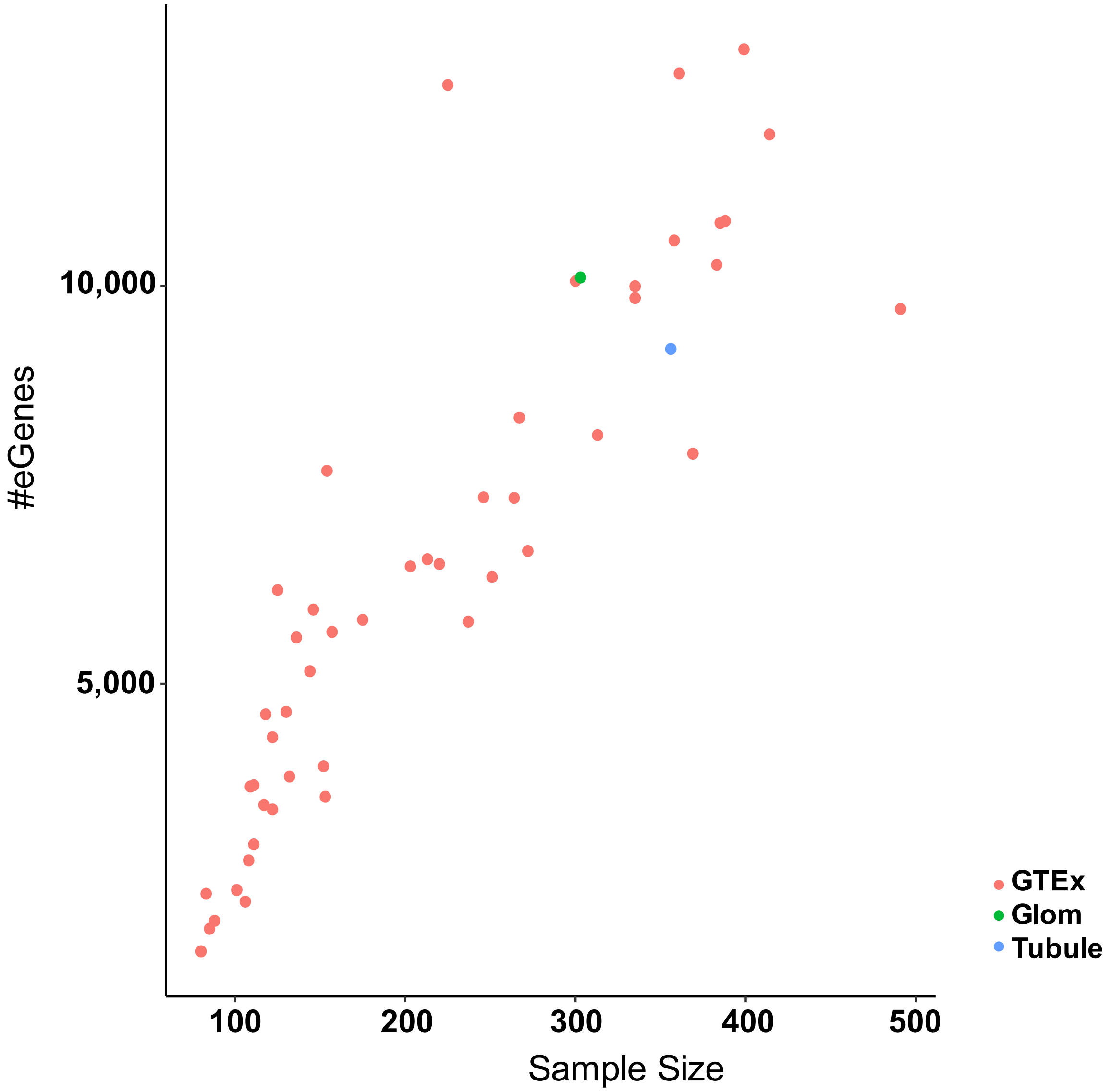


**Fig. S3. eQTLs identified in each kidney compartment.** Relationship between sample size (x axis) and the number of identified eGenes (y axis). Green data point, the number of eGenes identified in glomeruli; blue data point, the number of eGenes identified in tubuli; red dots, the number of eGenes identified 48 human tissue eQTL data from GTEx (v7).

**Supplementary figure 4**

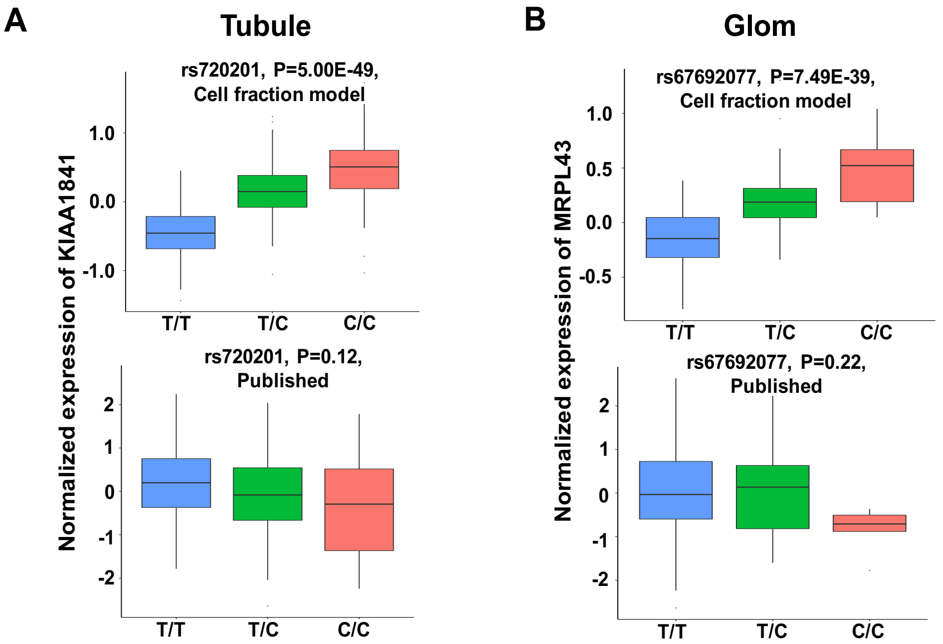


**Fig. S4. Example of newly identified eQTLs using eQTL(cf) model with large sample size for each kidney compartment comparing to our previous eQTL study (*1*).** A). Genotype of SNP rs720201 and the gene expression of *KIAA1841* association in human kidney tubules in new cell fraction adjusted eQTL (cf) data (top panel, N=356 samples) or the previously published tubule eQTL data (*1*) (bottom panel, N=121 samples). B). Genotype of SNP rs67692077 and the gene expression of *MRPL43* association in human kidney glomeruli in new cell fraction adjusted eQTL (cf) data (top panel, N=303 samples) or the previously published glomeruli eQTL data (*1*) (bottom panel, N=119 samples). Center lines show the medians; box limits indicate the $25^{th}$ and $75^{th}$ percentiles; whiskers extend to the $5^{th}$ and $95^{th}$ percentiles; outliers are represented by dots. P was calculated by linear regression model.

**Supplementary figure 5**

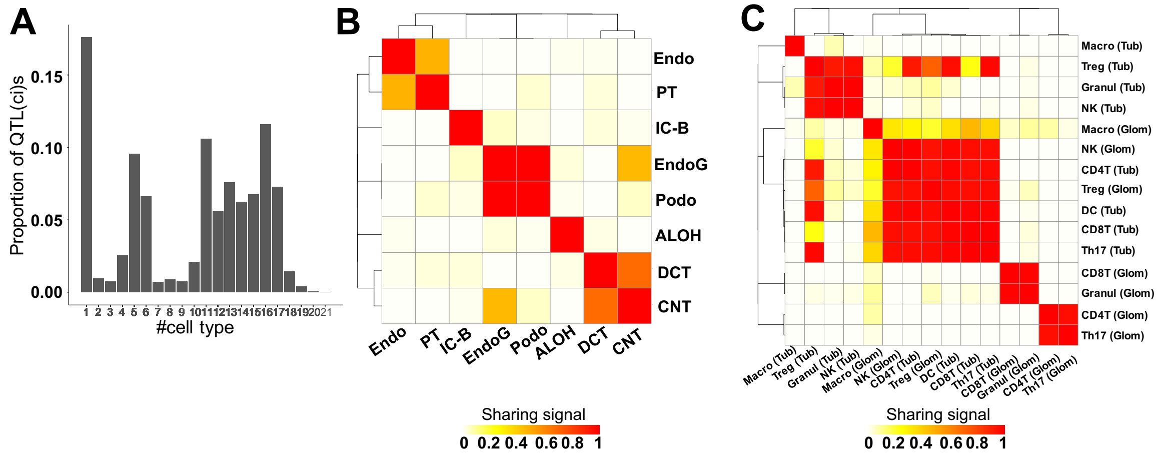


**Fig. S5. Sharing of eQTL(ci)s across cell types in tubuli and glomeruli.** A). About 18% eQTL(ci)s show cell type specificity across the 23 highly correlated cell types deconvoluted in each kidney compartment. Cell type specific eQTL(ci) was defined as the eQTL(ci) at LFSR<0.05 only in a single cell type. B). Pairwise sharing estimated by *mash* (*2*) of eQTL(ci)s across kidney cell types. C). Pairwise sharing estimated by *mash* (*2*) of eQTL(ci)s across immune cell types. The color-coded sharing signal is the proportion of significant eQTL(ci)s (LFSR<0.05) that are shared in the same sign and similar magnitude (effect within a factor of 0.5) between two cell types. EndoG: glomerular endothelial cells, Endo: endothelial cells, Podo: podocyte, PT: proximal tubule, ALOH: ascending loop of Henle, DCT: distal convoluted tubule, IC-B: beta intercalated cells, CNT: connecting tubule, CD4T: CD4 T cells, CD8T: CD8 T cells, DC: CD11b+ dendritic cells, Granul: granulocyte, Macro: macrophage, Th17: T helper 17 cells, Treg: regulatory T cells, NK: natural killer cells. Tub: tubule, Glom: glomeruli.

**Supplementary figure 6**

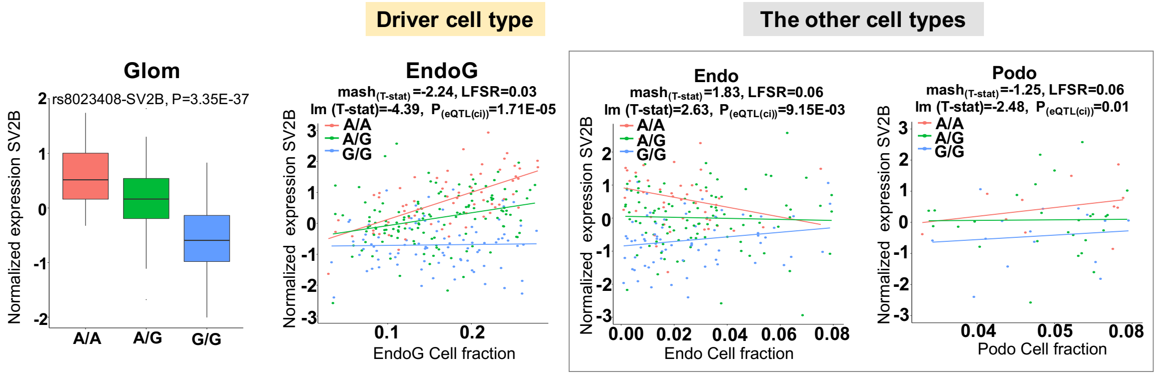


**Fig. S6. Defining cell-type dependent activities of genetic variants on gene expression.** EndoG specific eQTL(ci) at the *SV2B* gene. The leftmost panel illustrates the association between genotype of SNP rs8023408 and gene expression of *SV2B* in human glomeruli (N=303 samples). The y-axis is the normalized expression of *SV2B* in human kidney glomeruli, the x-axis is the genotype at rs8023408 locus. Center lines show the medians; box limits indicate the 25$^{th}$ and 75$^{th}$ percentiles; whiskers extend to the 5$^{th}$ and 95$^{th}$ percentiles; outliers are represented by dots. P value was calculated by linear regression eQTL(cf) model. The following panels show EndoG cell-type-specificity of this eQTL(ci) across 3 different kidney function cell types in human glomeruli (N=303 samples). The y-axis is the normalized expression of *SV2B* in human kidney glomeruli (N=303 samples), the x-axis is the cell fraction for each specific kidney function cell type in subjects with genotype A/A (red), A/G (green) and G/G (blue) at rs8023408 locus. lm (T-stat): T-statistics calculated from linear model of eQTL (ci), mash$_{(T\text{-stat})}$: T-statisitcs estimated by *mash*. EndoG: glomerular endothelial cell, Endo: endothelial, Podo: podocyte. Each data point represents for an individual.

**Supplementary figure 7**

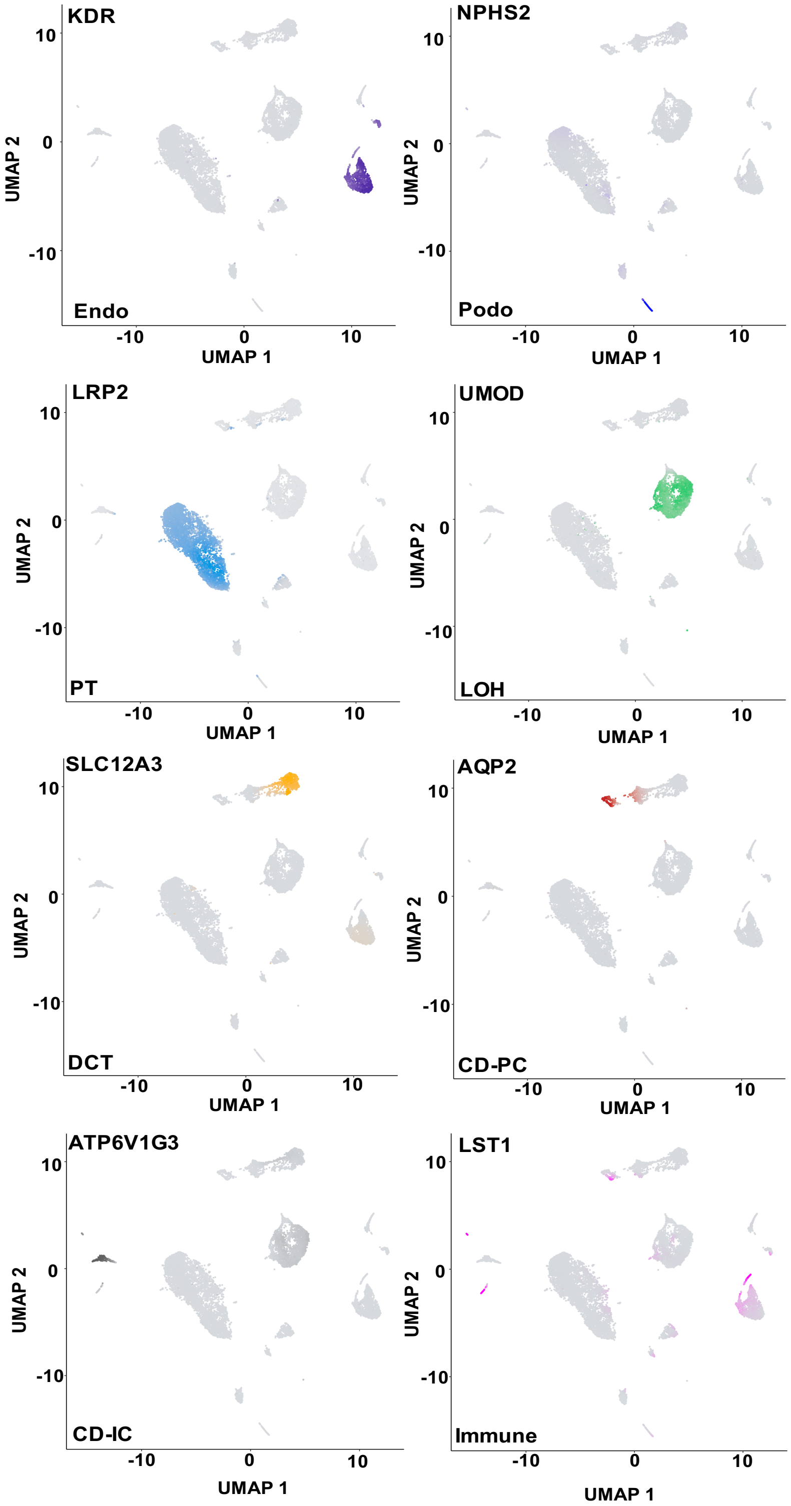


**Fig. S7. Feature plots of open chromatin information for known cell-type specific marker genes (*3, 4*).**

**Supplementary figure 8**

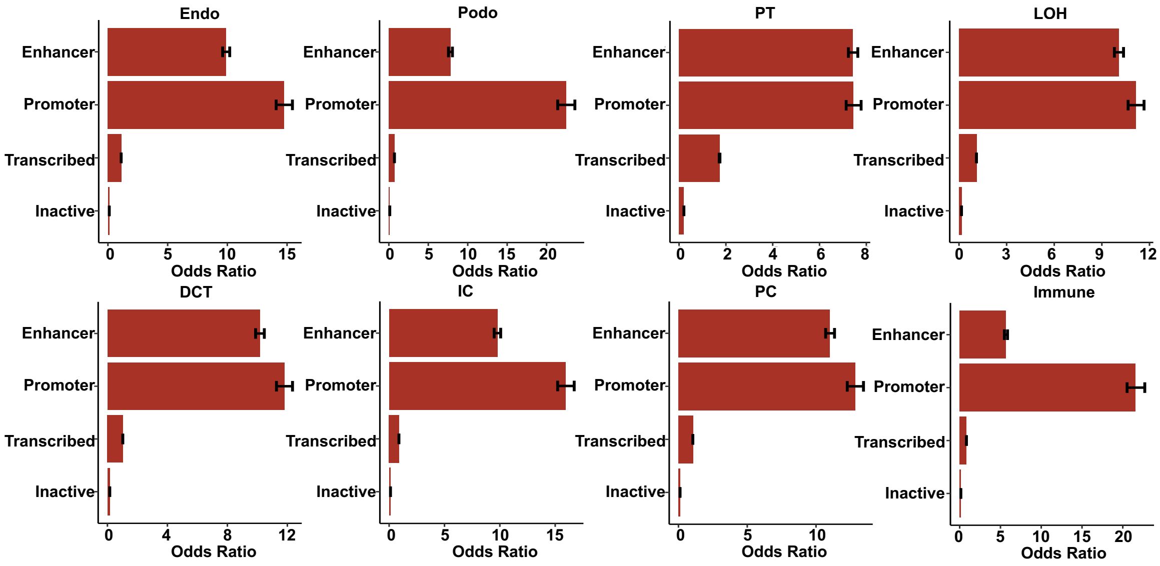


**Fig. S8. The enrichment (Odds Ratio, X-axis) of open chromatin peaks for each cell type called from the snATAC-seq data in the different human kidney chromatin states (Y-axis).** The error bars denote the 95% confidence intervals of Odds Ratio. Endo: endothelial cells, Podo: podocyte, PT: proximal tubule, LOH: loop of Henle, DCT: distal convoluted tubule, PC: collecting duct principal cells, IC: collecting duct intercalated cells, Immune: Immune cells.

**Supplementary figure 9**

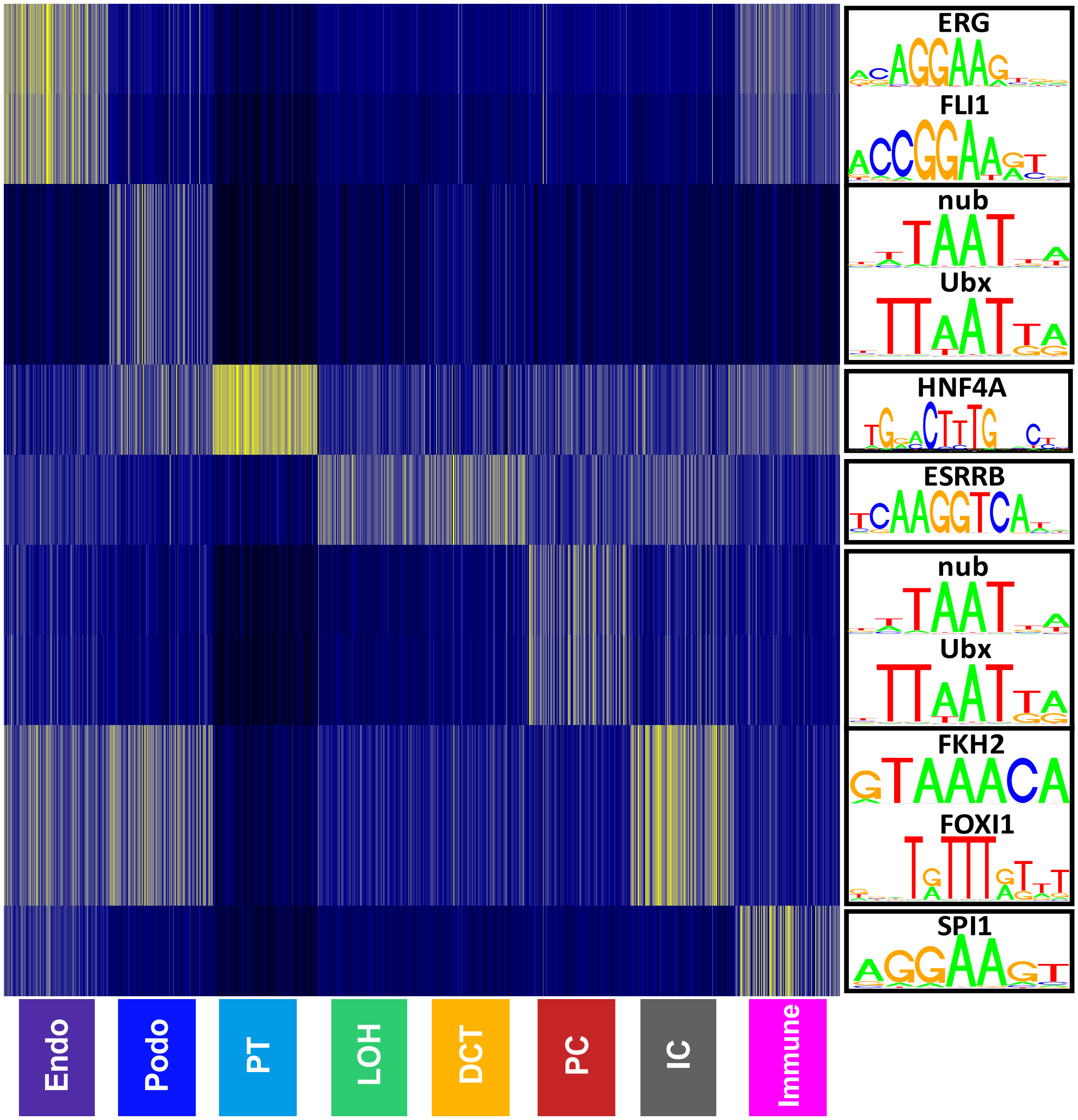


**Fig. S9. Motif enrichment analysis using cell type specific differential accessibility peaks (DAPs)** Each row represents the top enriched motif for each cell type. Each column represents each cell. For each cell type, we randomly selected 100 cells for each cell type to show the cell type specific motif enrichment. Endo: endothelial cells, Podo: podocyte, PT: proximal tubule, LOH: loop of Henle, DCT: distal convoluted tubule, PC: collecting duct principal cells, IC: collecting duct intercalated cells, Immune: Immune cells.

**Supplementary figure 10**

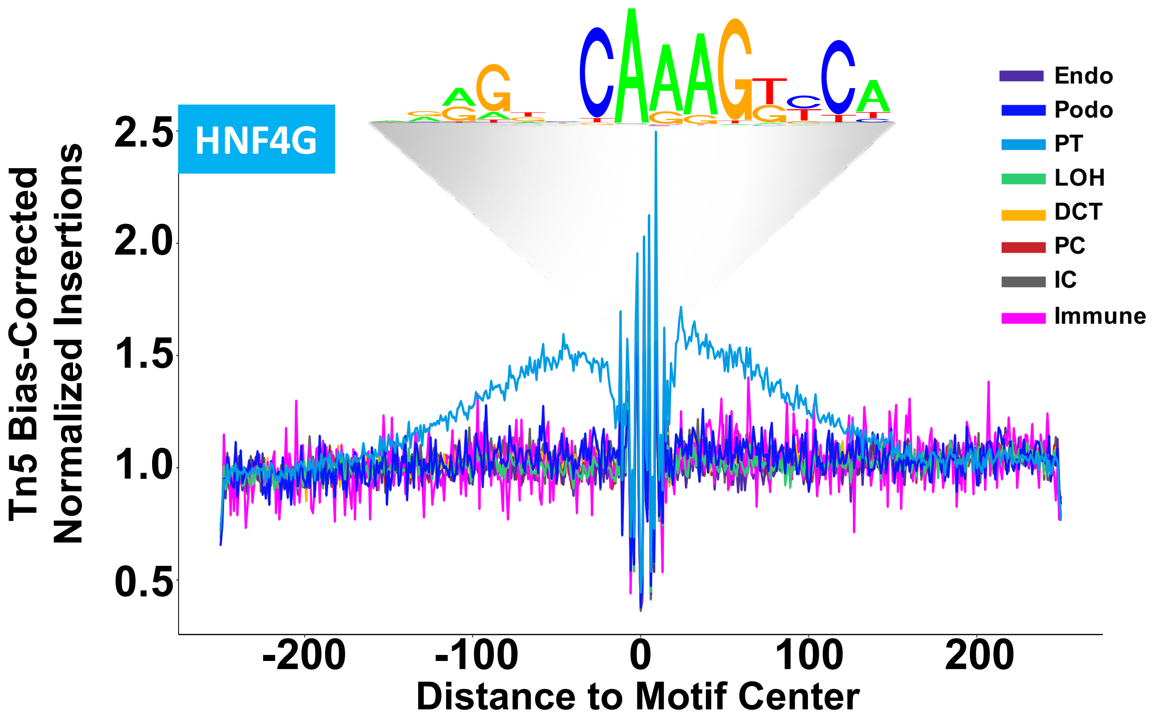


**Fig. S10. Footprinting analysis of the HNF4G transcription factor across the 8 major cell types.** The motif logos are shown above. Endo: endothelial cells, Podo: podocyte, PT: proximal tubule, LOH: loop of Henle, DCT: distal convoluted tubule, PC: collecting duct principal cells, IC: collecting duct intercalated cells, Immune: Immune cells.

**Supplementary figure 11**

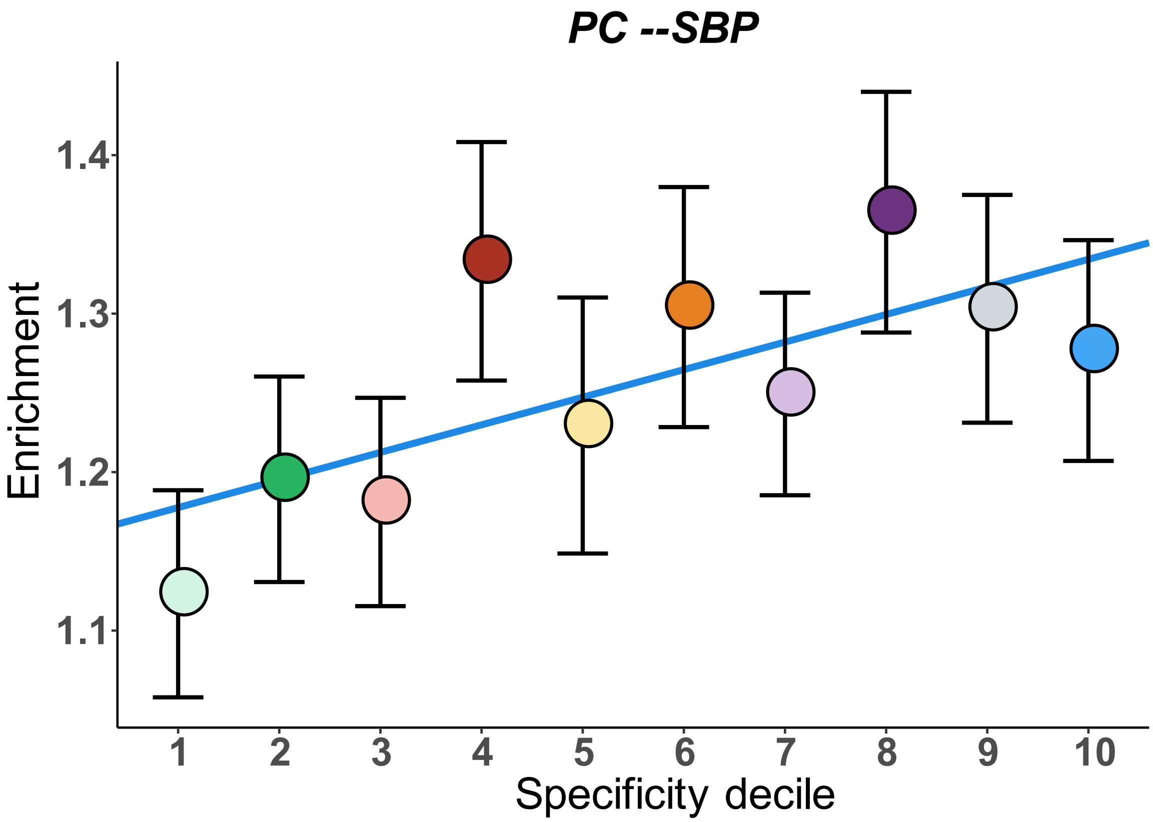


**Fig. S11. Enrichment of SBP-SNP heritability in each of the specificity deciles for collecting duct principal (PC) cells** calculated using LDSC from kidney scRNA-seq data (*3*)**.** X-axis is the gene expression specificity decile, Y-axis is the enrichment value calculated by LDSC. Colors of dots represent different specificity deciles. Blue line shows the linear regression slope fitted to the enrichment values. Error bars indicate the 95% confidence intervals.

**Supplementary figure 12**

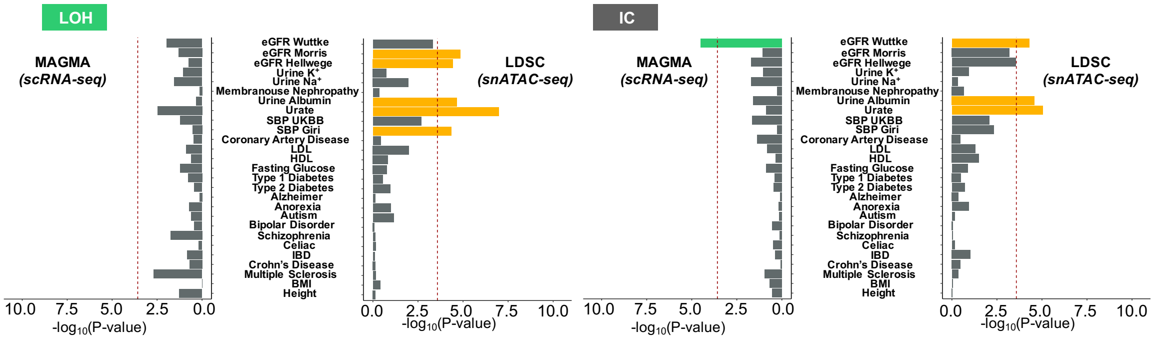


**Fig. S12. Evaluation of enrichment of common GWAS variants in each kidney cell type from mice kidney scRNA-seq data(*3*) using MAGMA (the left panels) and human kidney snATAC-seq data using LDSC (the right panels).** X-axis represents 24 different GWAS traits and y-axis is the $-\log_{10}$(P-value). P-value was estimated by MAGMA (the left panels) and LDSC (the right panels). LOH: loop of Henle, IC: collecting duct intercalated cells.

### Supplementary figure 13

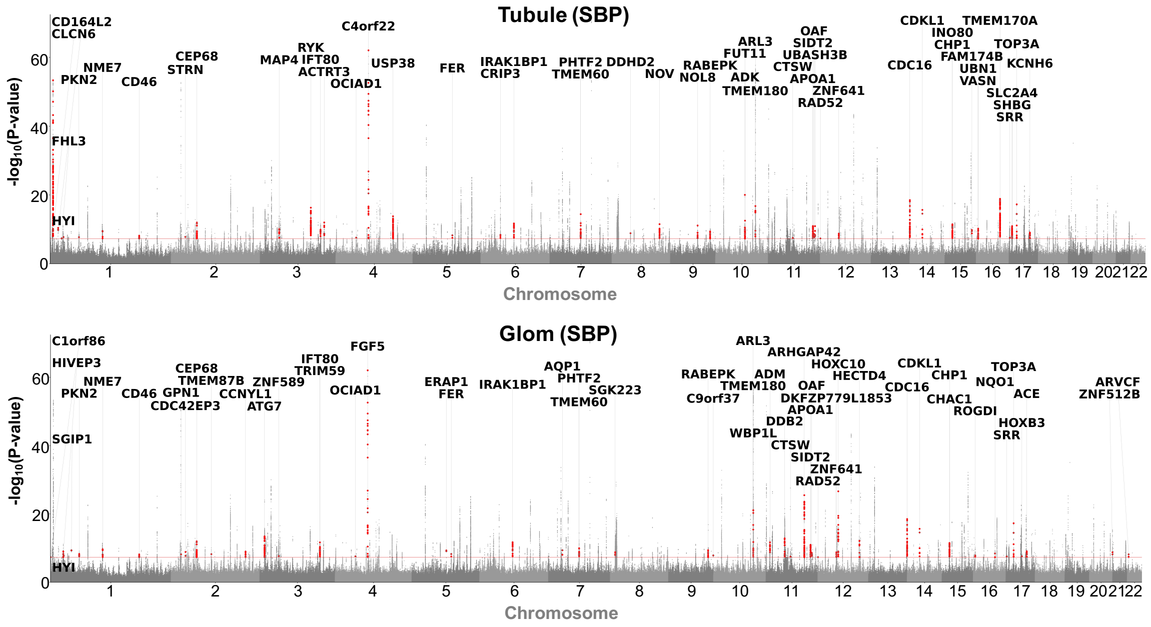


**Fig. S13. Miami plot showing coloc nominated target genes for the SBP GWAS associated loci identified by eQTL(cf) model in tubules and glomeruli across the whole genome.** X-axis represents the chromosome. Y-axis represents the GWAS significance –log$_{10}$(P-value).

**Supplementary figure 14**

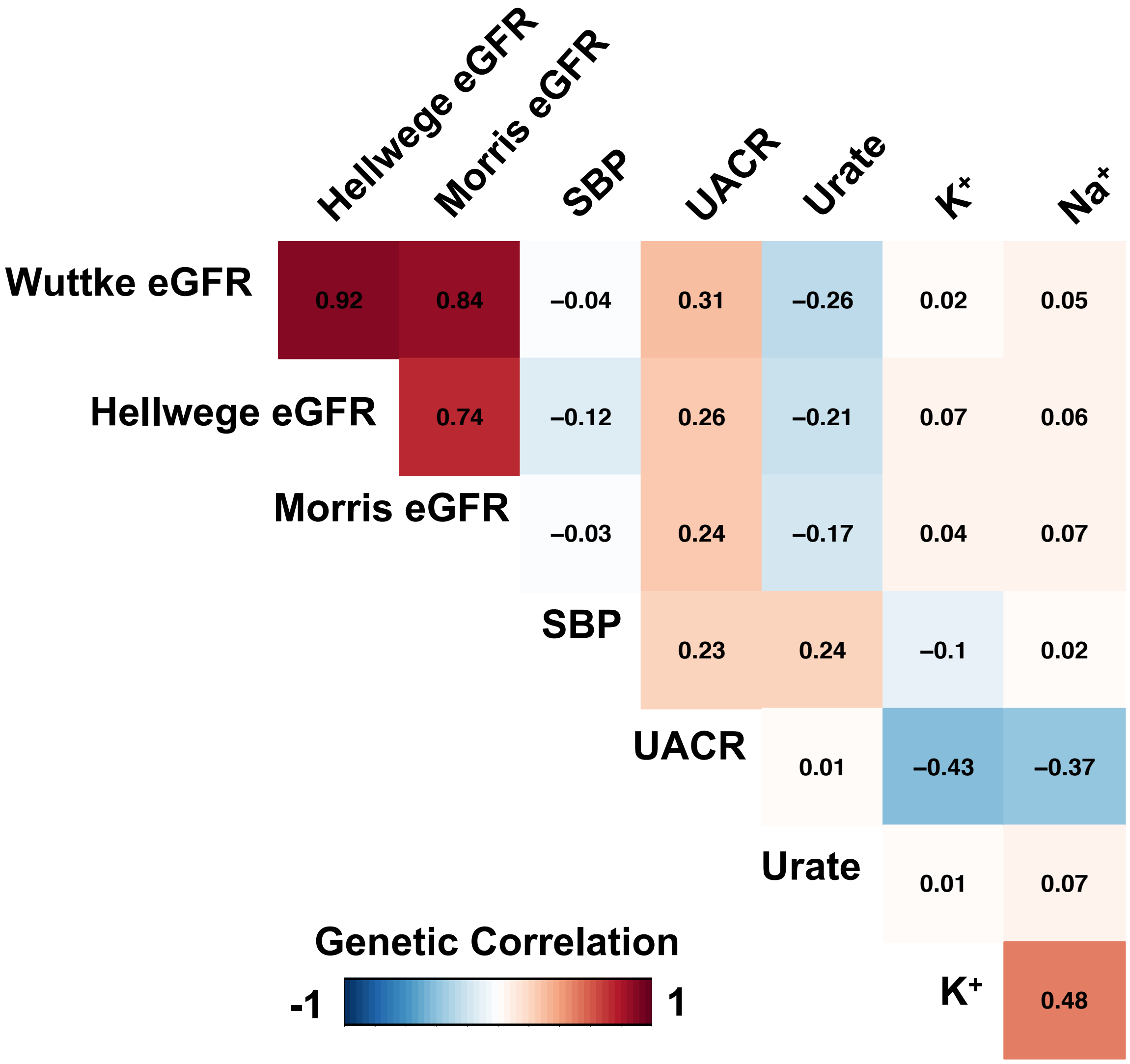


**Fig. S14. Pairwise genetic correlation of 6 kidney related GWAS endophenotypes**

**Supplementary figure 15**

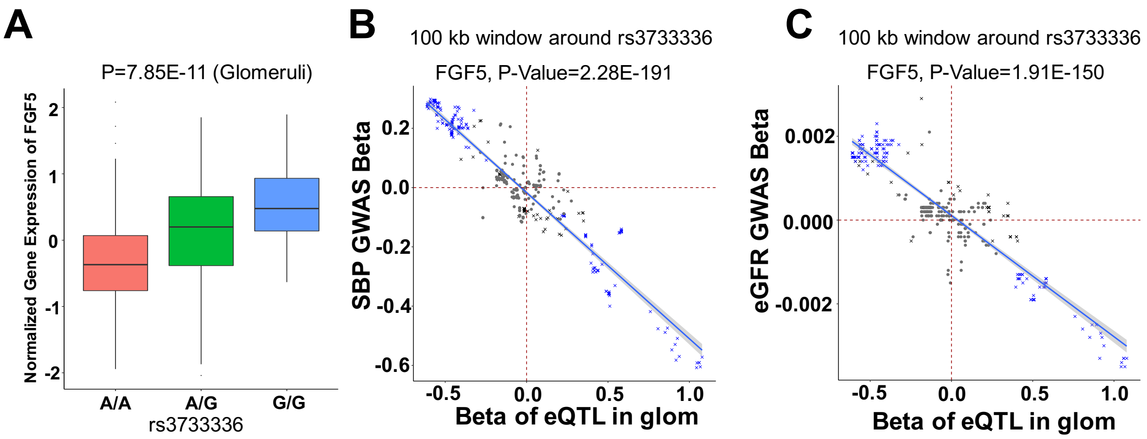


**Fig. S15. *FGF5* gene prioritization for kidney function and blood pressure.** A). The association of genetic variant rs3733336 and gene expression of *FGF5* in 303 human glomerular samples. P value was calculated by linear regression eQTL (cf) model. Center lines show the medians; box limits indicate the 25$^{th}$ and 75$^{th}$ percentiles; whiskers extend to the 5$^{th}$ and 95$^{th}$ percentiles; outliers are represented by dots. P was calculated by linear regression model.

B). The variant effect sizes of SBP GWAS and *FGF5* expression changes (eQTL(cf)), located within ±100 kb of rs3733336 are significantly correlated. Each data point represents a single SNP. If the P-values of eQTL or GWAS were <0.05, the data points are shown as a 'cross', if the P-values of both eQTL and GWAS were <0.05, the data points are shown in 'blue'. The sample size is N=303 for eQTL in glomeruli; N=477,054 samples for SBP GWAS.

C). The variant effect sizes eGFR GWAS, and *FGF5* expression changes (eQTL(cf)), located within ±100 kb of rs3733336 are significantly correlated. Each data point represents a single SNP. If the P-values of eQTL or GWAS were <0.05, the data points are shown as a 'cross', if the P-values of both eQTL and GWAS were <0.05, the data points are shown in 'blue'. The sample size is N=303 for eQTL in glomeruli; N=765,348 samples for eGFR GWAS.

**Supplementary figure 16**

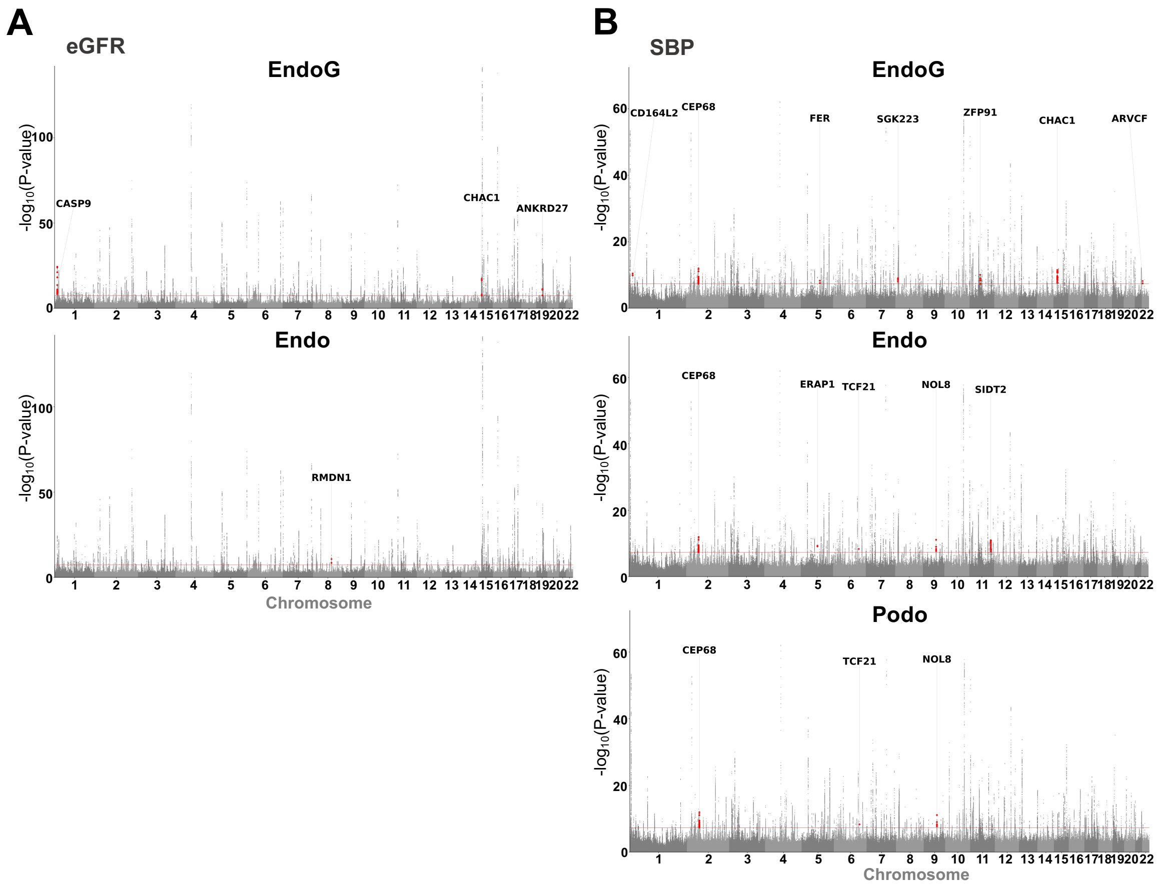


**Fig. S16. Miami plot showing eQTL(ci) coloc prioritized eGFR (A) and SBP (B) GWAS associated genes.** X-axis represents the chromosomal location. Y-axis represents the GWAS significance $-\log_{10}$(P-value). EndoG: glomerular endothelial cell, Endo: endothelial cells, Podo: podocyte.

**Supplementary figure 17**

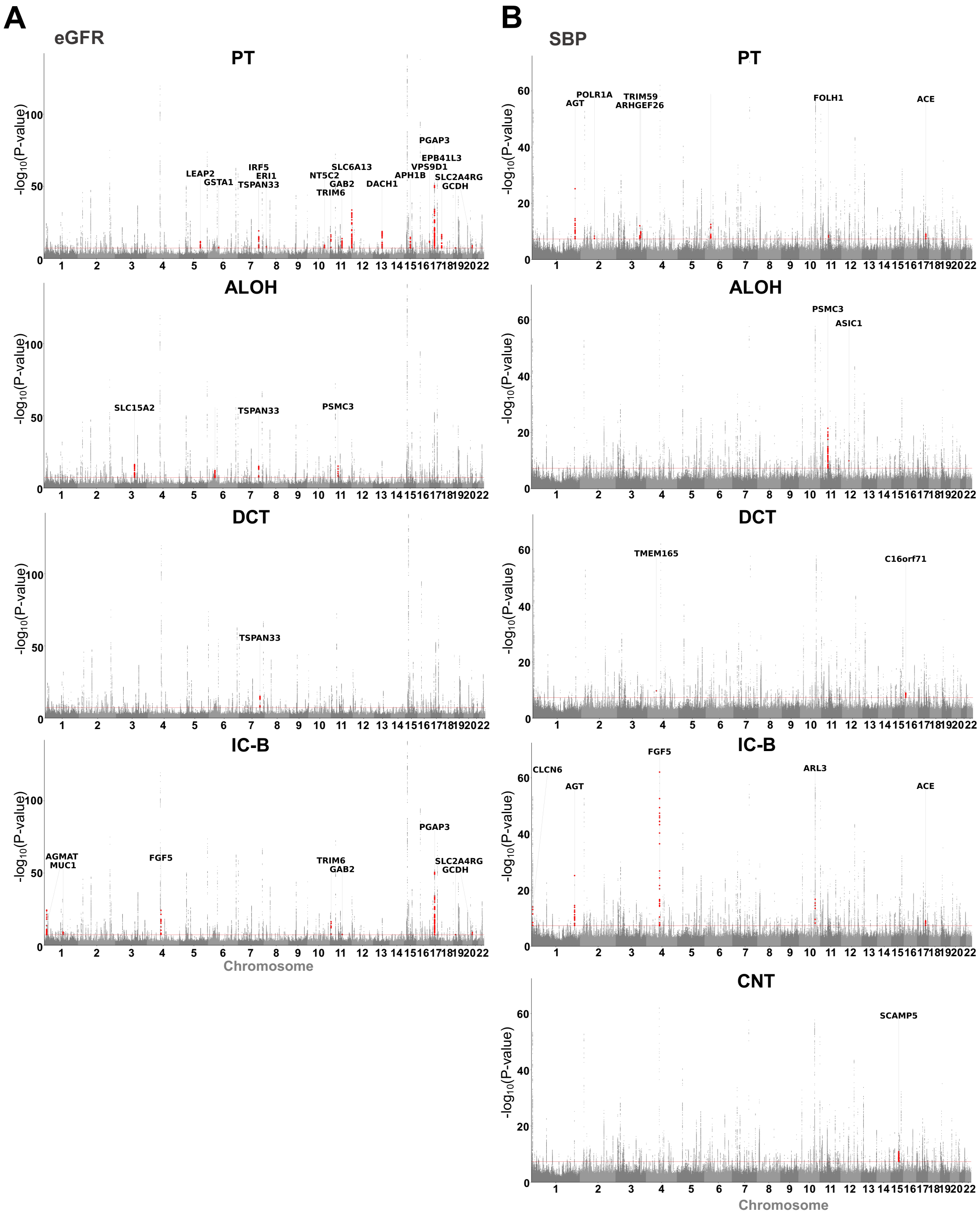


**Fig. S17. Miami plot showing eQTL(ci) coloc prioritized genes for eGFR (A) and SBP (B) GWAS.** X-axis represents the chromosomal location. Y-axis represents the GWAS significance –$\log_{10}$(P-value). PT: proximal tubule, ALOH: ascending loop of Henle, DCT: distal convoluted tubule, IC-B: beta intercalated cells, CNT: connecting tubule.

**Supplementary figure 18**

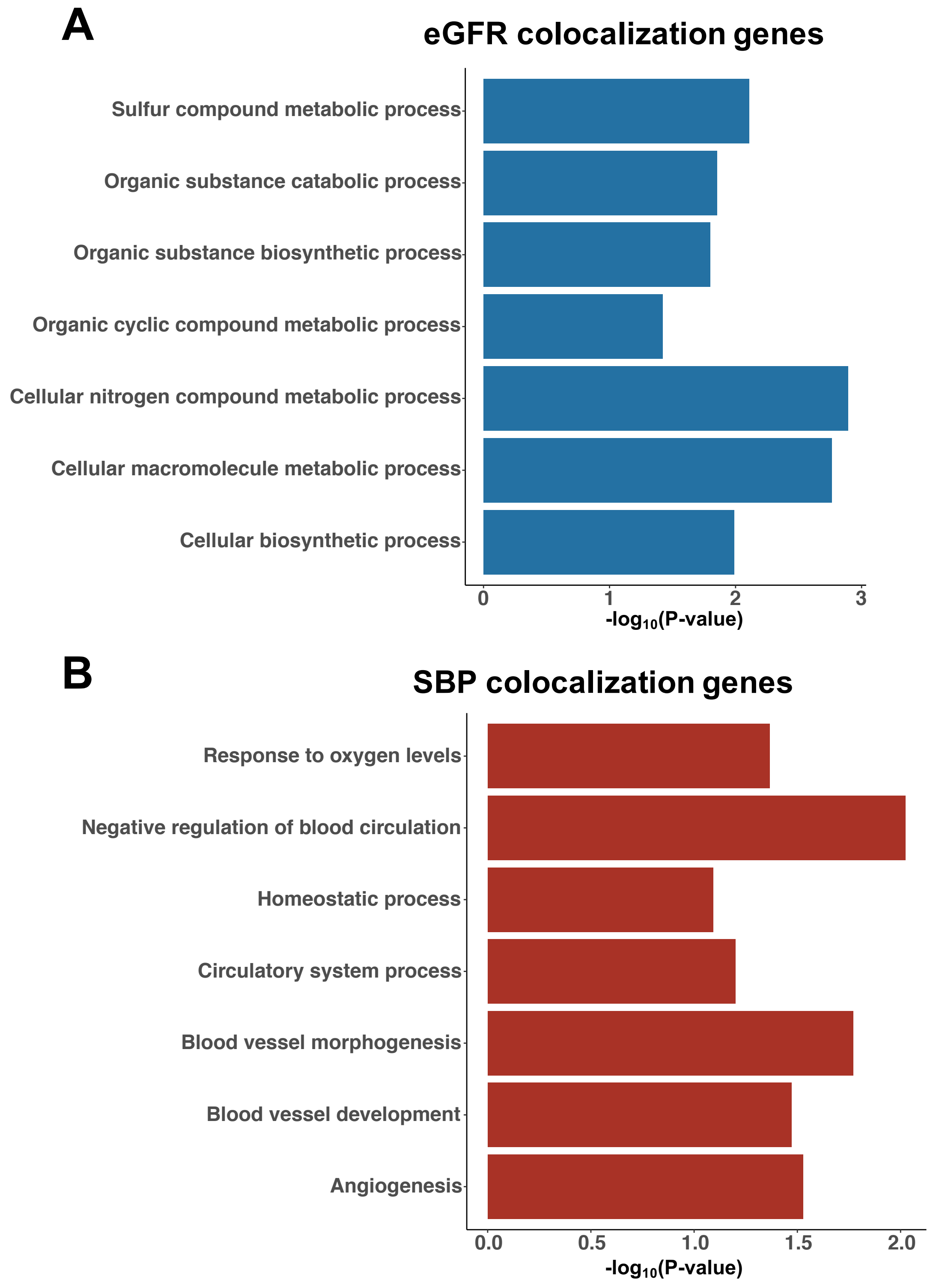


**Fig. S18. Gene ontology based functional annotation of eGFR (A) and SBP (B) GWAS and eQTL colocalization nominated genes (DAVID).**

**Supplementary figure 19**

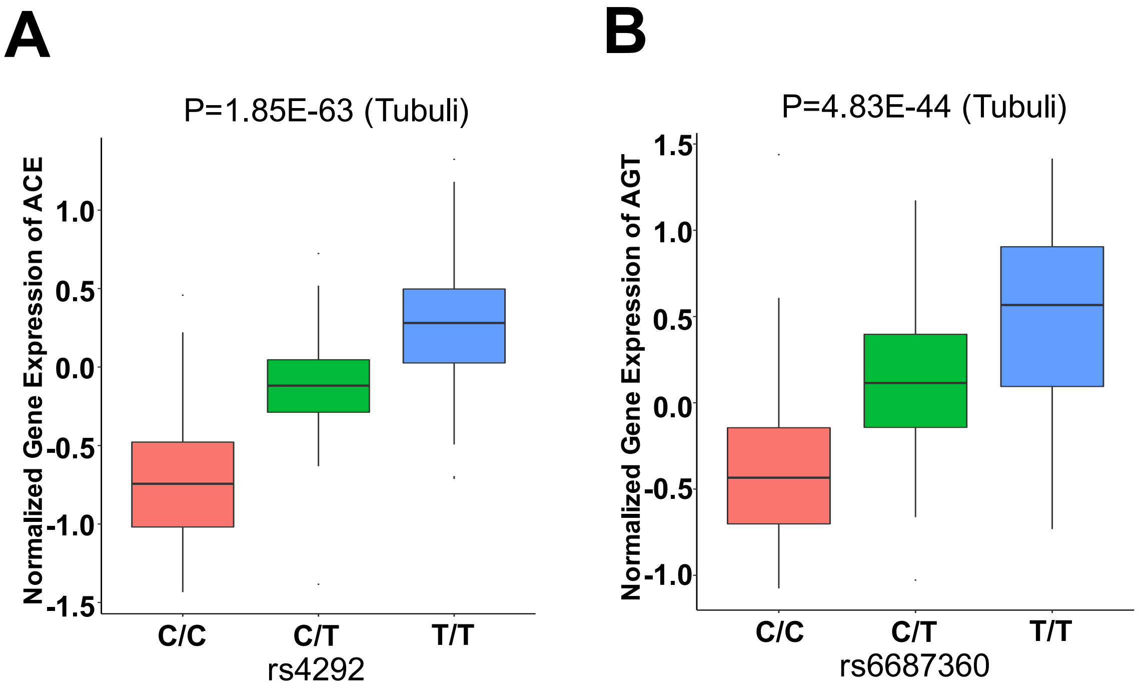


**Fig. S19. Human kidney tubule eQTL(cf)s of rs4292-*ACE* and rs6687360-*AGT*.** A. The association of genetic variant rs4292 and gene expression of *ACE* in 356 human tubuli samples. B. The association of genetic variant rs6687360 and gene expression of *AGT* in 356 human tubuli samples. P value was calculated by linear regression eQTL (cf) model. Center lines show the medians; box limits indicate the 25$^{th}$ and 75$^{th}$ percentiles; whiskers extend to the 5$^{th}$ and 95$^{th}$ percentiles; outliers are represented by dots. P was calculated by linear regression model.

### Supplementary figure 20

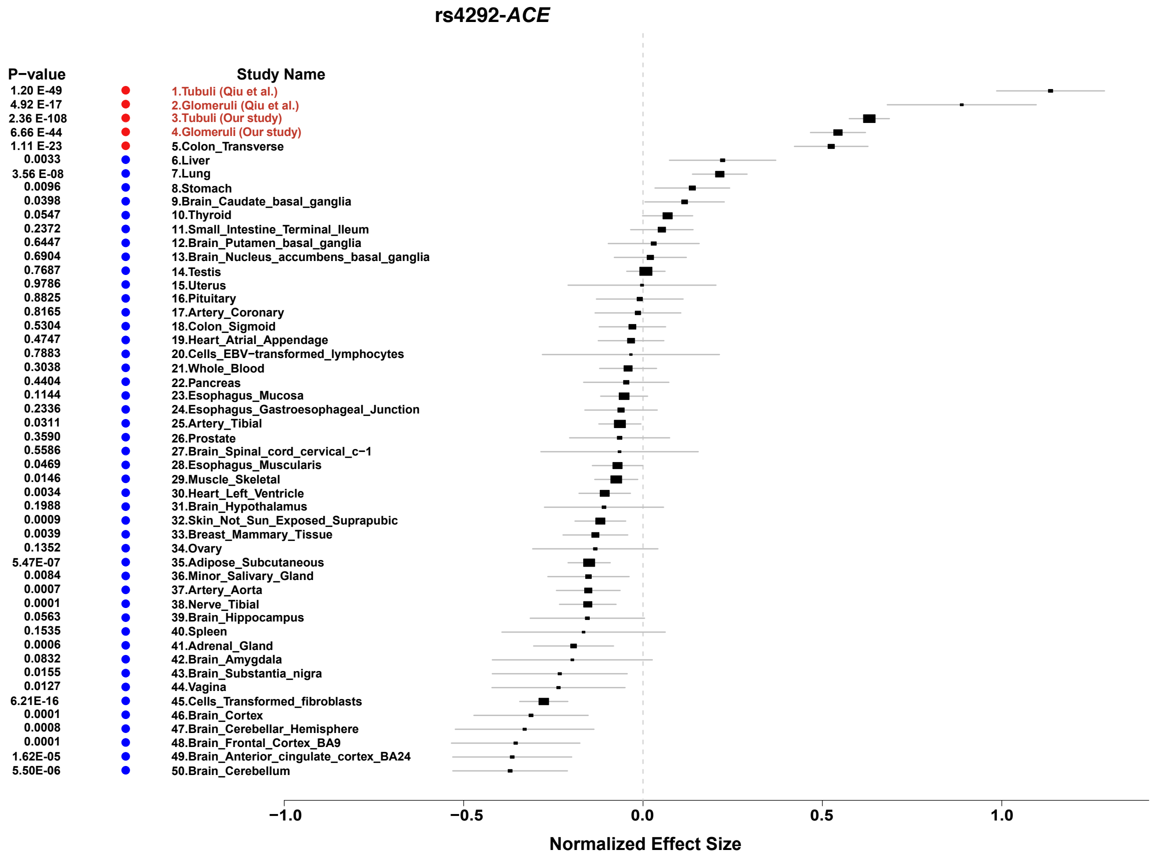


**Fig. S20. Meta-analysis of eQTL rs4292-*ACE* in kidney compartments and 46 GTEx tissues.** The normalized effect size (**β**) of eQTL association between rs4292 and *ACE* in tubular (n=356 for this study and n=121 for study of Qiu et al.(*1*)) and glomerular (n=303 and n=121 for glomerular of Qiu et al.) compartments and in 46 human tissues from GTEx (v7). The x axis shows **β** with the 95% confidence interval (CI). EBV, Epstein-Barr virus.

**Supplementary figure 21**

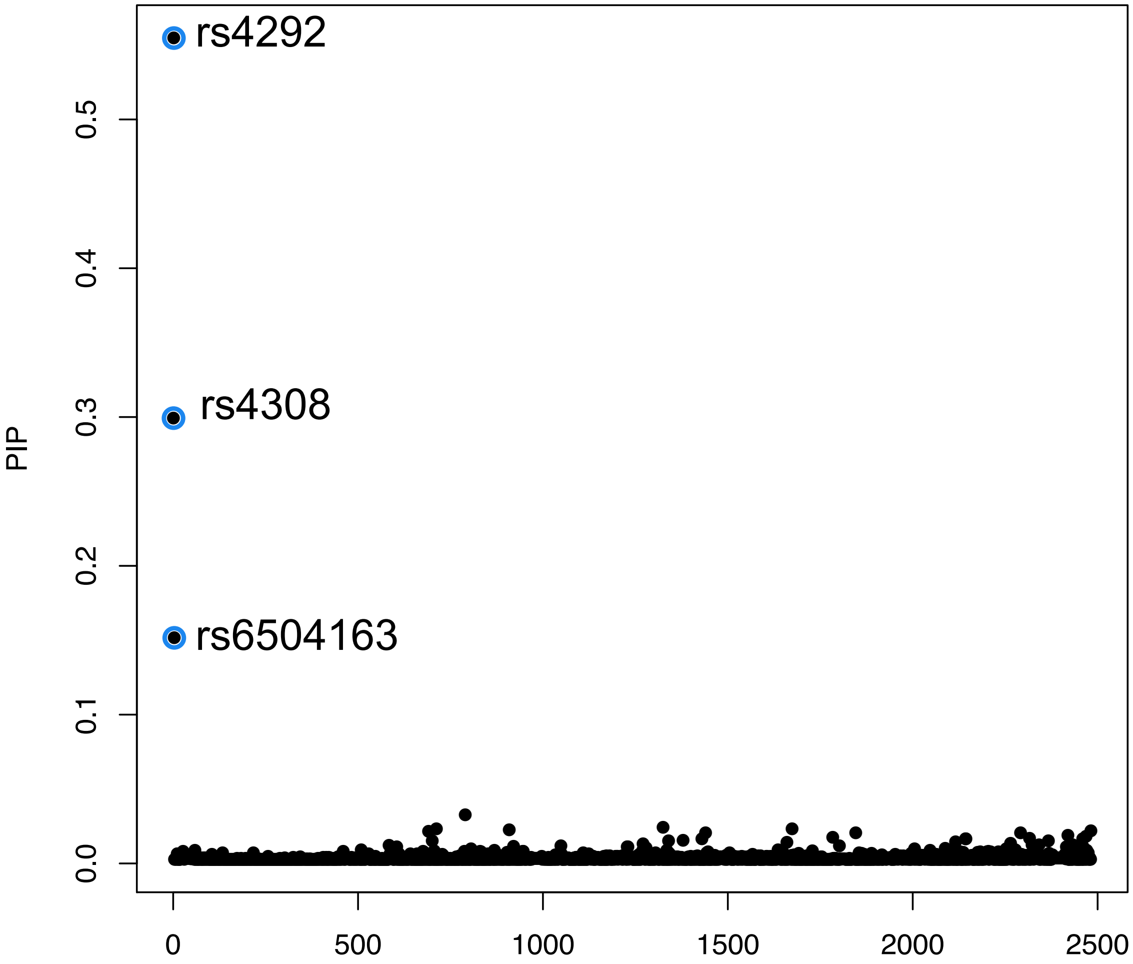


**Fig. S21. Fine mapping results of causal variants affecting *ACE* gene expression in human kidney tubule samples.** There are about 2,500 variants (X-axis) locate in the ± 100 Mb region around the transcription start site (TSS) of ACE. Only three SNPs were identified as the potential causal variants that driving the gene expression changes in tubuli. rs4292 is the top variant with the highest posterior probability (Y-axis) (PIP=55.5%).

**Supplementary figure 22**

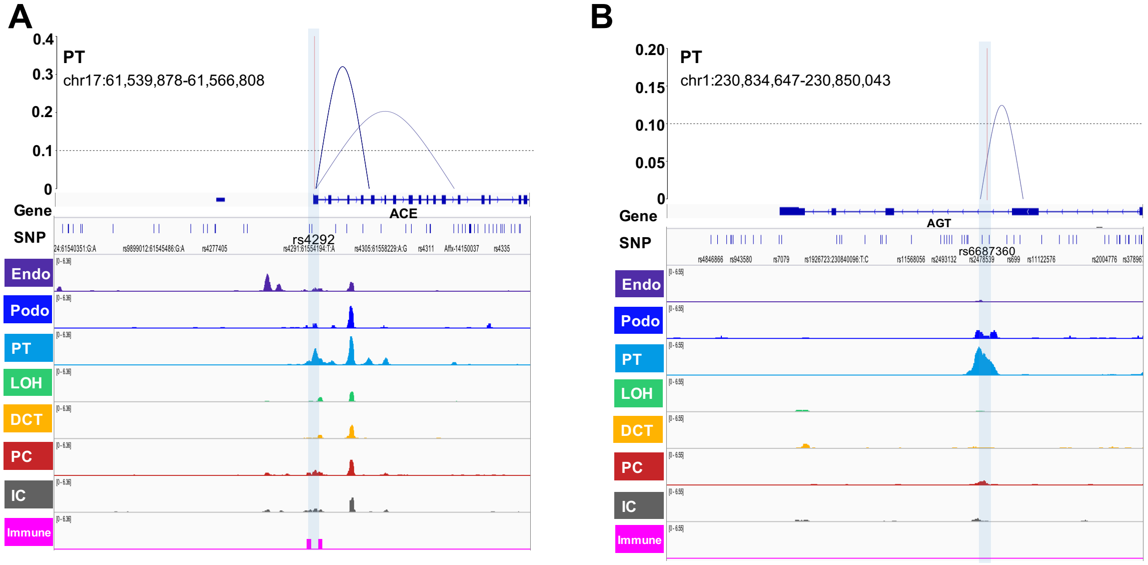


**Fig. S22. Cicero-inferred co-accessibility between pairs of accessible chromatin sites at the A). *ACE* promoter and B). *AGT* gene body region.** The showing genomic region are A). chr17:61,539,878-61,566,808 and B). chr1:230,834,647-230,850,043.

**Supplementary figure 23**

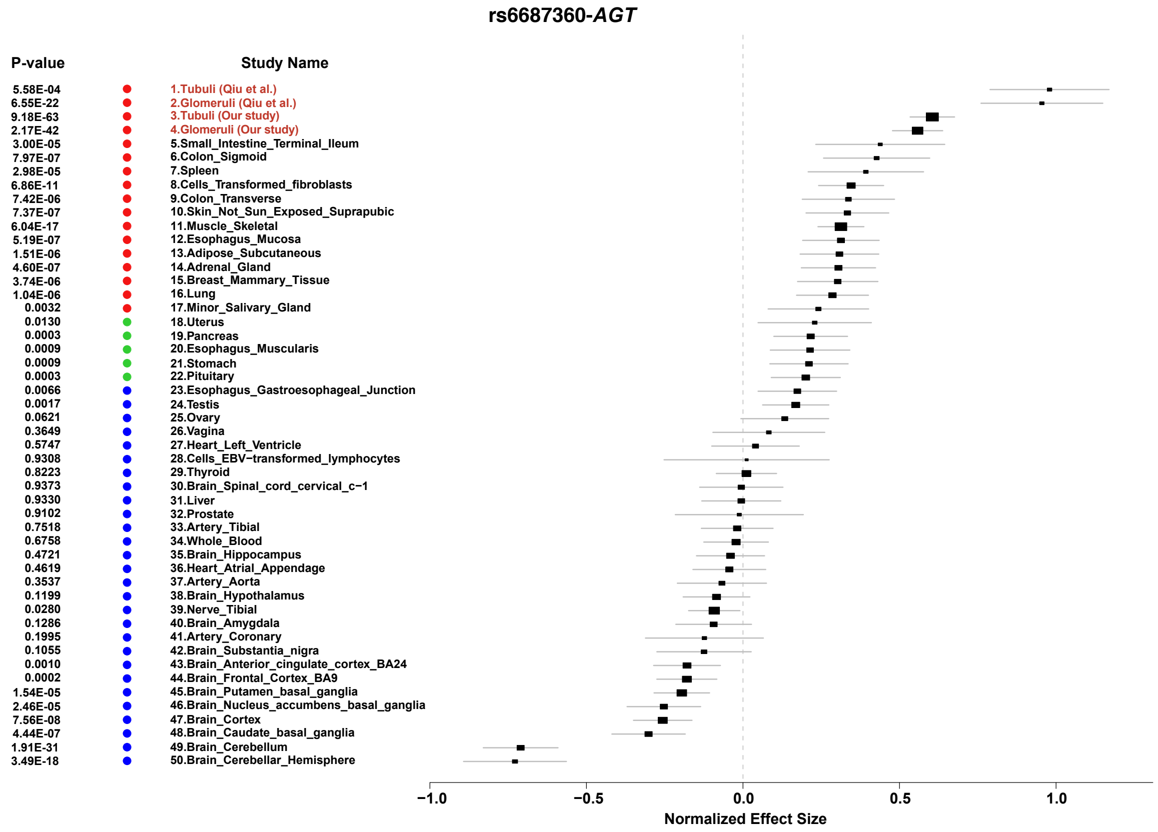


**Fig. S23. Meta-analysis of eQTL rs6687360-*AGT* in human kidney compartments and 46 GTEx tissues.** The normalized effect size (**β**) of eQTL association between rs6687360 and *AGT* in tubular (n=356 for this study and n=121 for study of Qiu et al.(*1*)) and glomerular (n=303 and n=121 for glomerular of Qiu et al.) compartments and in 46 human tissues from GTEx (v7). The x axis shows **β** with the 95% confidence interval (CI). EBV, Epstein-Barr virus.

**Supplementary figure 24**

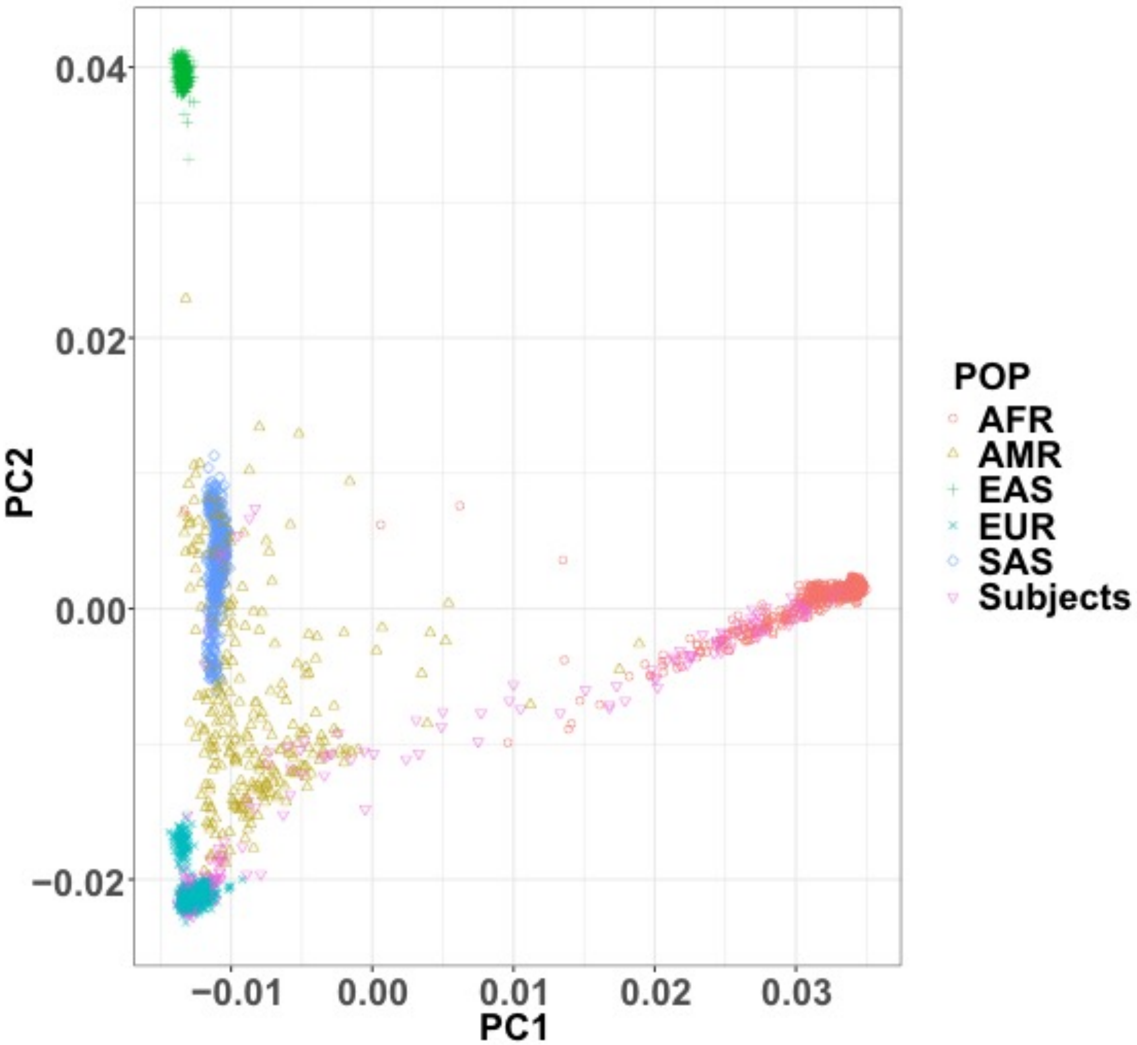


**Fig. S24. Genotype data distribution** Principal component analysis of genotype data of 367 participants (individuals with high quality genotype data available) and 1000 Genomes Project Phase 3 reference genome data. Magenta triangles indicate 367 kidney samples (of the current study).

**Supplementary figure 25**

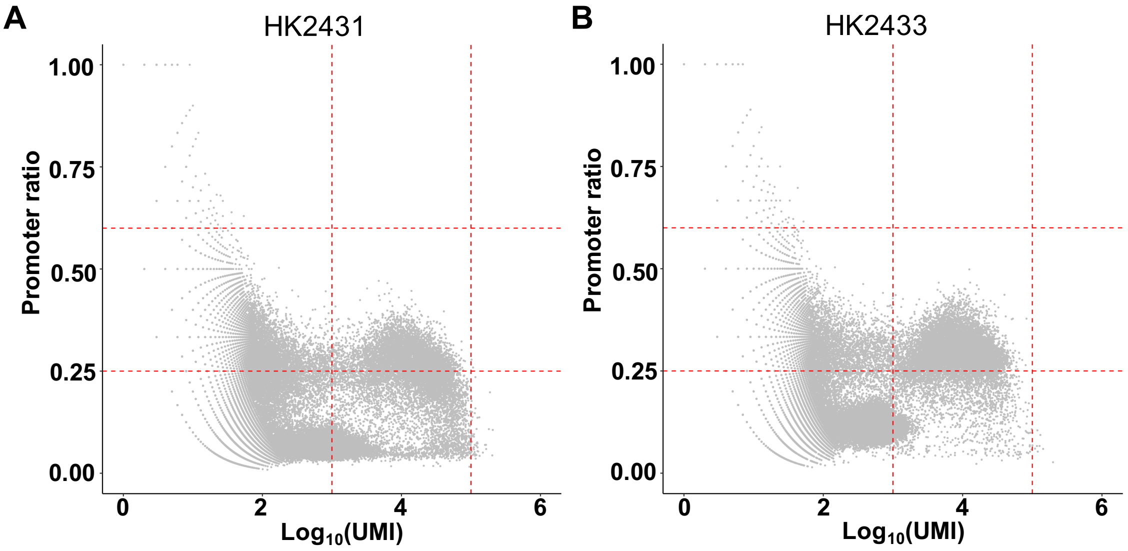


**Fig. S25. Quality Control of human kidney snATAC-seq data** Each data point represents a cell, x-axis: $\log_{10}$(UMI), UMI: unique molecular identifier; y-axis: promoter ratio of sample A. HK2431 B. HK2433. Only cells with logUMI in [3-5] (red dash lines) and promoter ratio in [0.25-0.6] (red dash lines) were retained.

**Supplementary figure 26**

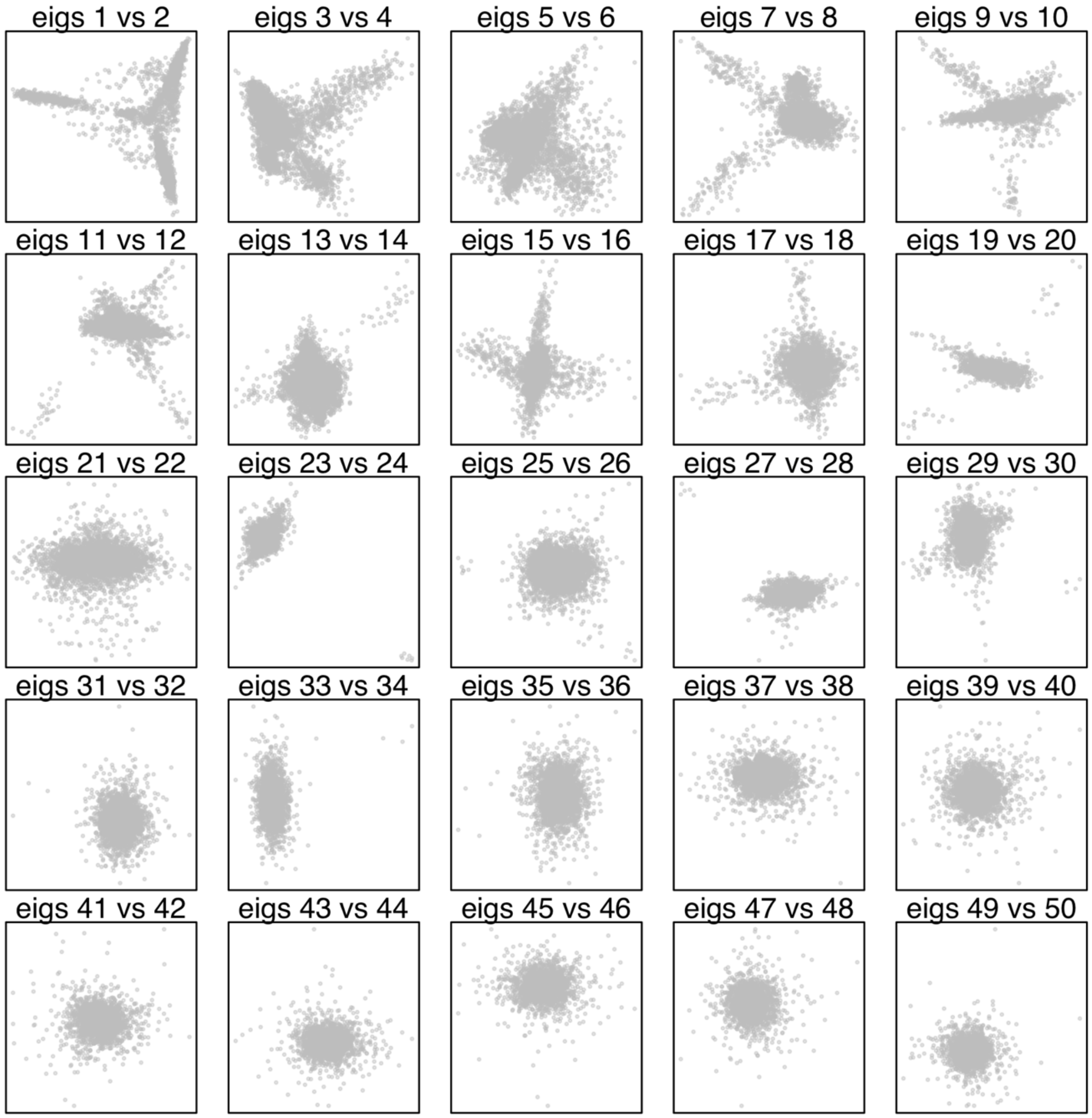


**Fig. S26**. **Human kidney snATACseq dimensionality reduction** 50 eigenvectors generated by function *runDiffusionMaps* of the SnapATAC (*5*) package.

**Supplementary figure 27**

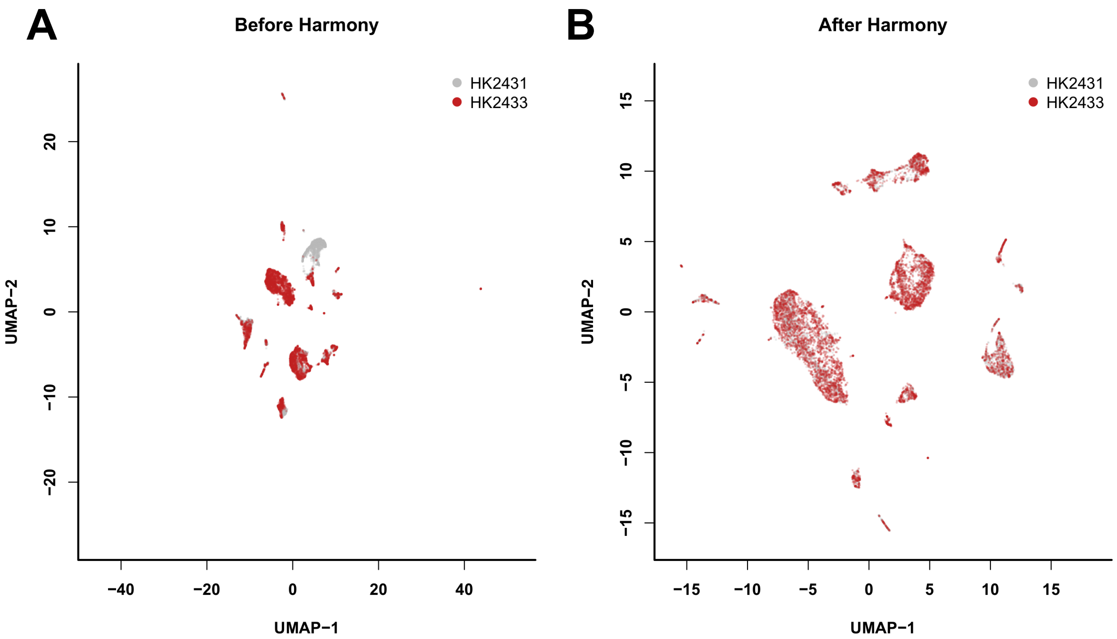


**Fig. S27. Batch effect correction of human kidney snATACSeq by Harmony** (*6*) A) Before batch effect correction B) After batch effect correction.

**Supplementary figure 28**

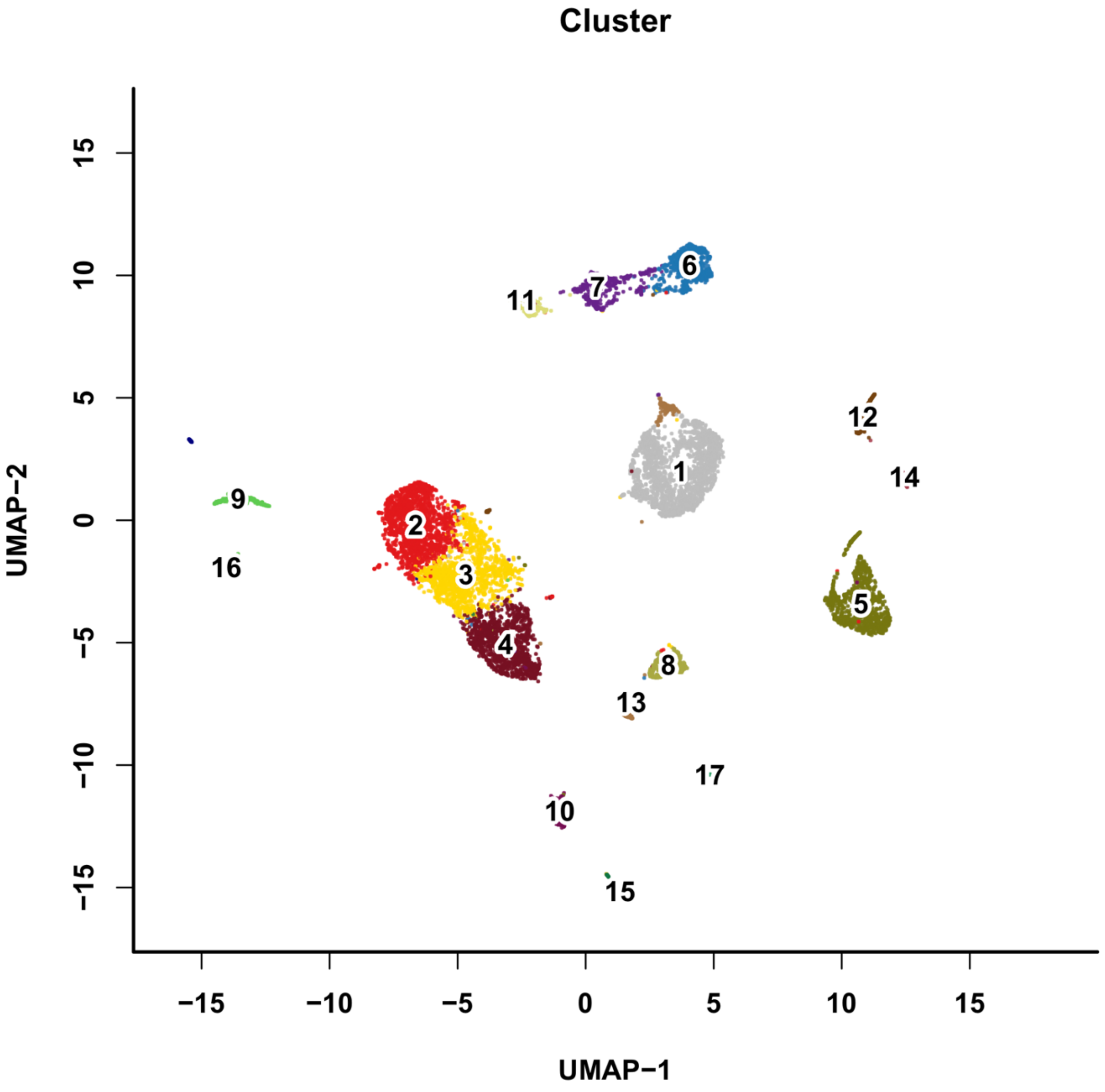


**Fig. S28. The initial UMAP of the human kidney snATAC-seq**
